# The Joubert syndrome protein CSPP1 is a conserved regulator of vertebrate multiciliogenesis and motile cilia function

**DOI:** 10.64898/2026.03.20.713242

**Authors:** Irem S. Dilbaz-Gunden, Clothilde Boitel, Jovana Deretic, Marine Touret, Mehmet S. Aydin, Esra Yigit, Ozgecan Kayalar, Hasan Bayram, Virginie Thomé, Olivier Rosnet, Nicolas Brouilly, Laurent Kodjabachian, Camille Boutin, Elif Nur Firat-Karalar

**Affiliations:** Department of Molecular Biology and Genetics, Koç University, Istanbul, Turkey 34450; Aix Marseille Univ, CNRS, IBDM, Turing Centre for Living Systems, Marseille, France; Mediopol University, Turkey; School of Medicine, Koç University, Istanbul, Turkey 34450

**Keywords:** CSPP1, cilia, basal bodies, centriole amplification, multiciliated cells, mouse tracheal epithelial cells, *Xenopus* epidermis

## Abstract

Cilia are conserved microtubule-based organelles required for signaling and fluid transport, and their dysfunction causes ciliopathies. Clinical overlap between sensory and motile ciliopathies suggests that primary and motile ciliogenesis depend on shared regulatory modules. Here, we identify Centrosome and Spindle Pole-associated Protein 1 (CSPP1), a microtubule-associated protein mutated in the neurodevelopmental ciliopathy Joubert syndrome, as a conserved regulator of vertebrate multiciliogenesis. Using mouse tracheal epithelial cultures and *Xenopus* embryonic epidermis, we show that CSPP1 localizes to fibrous granules and deuterosomes during centriole amplification, and to basal bodies and ciliary axonemes in differentiated multiciliated cells. Loss of CSPP1 impairs centriole amplification, basal body apical migration, spacing, and rotational polarity, and is accompanied by disorganization of the apical microtubule network. CSPP1 depletion also disrupts axoneme assembly, resulting in fewer and shorter cilia with ultrastructural defects, reduced ciliary beating, and impaired cilia driven fluid flow *in vivo*. Together, our findings identify CSPP1 as a conserved regulator of multiciliogenesis and motile cilia function and establish a basis for future work on how shared cytoskeletal pathways may underlie overlapping features of sensory and motile ciliopathies.

## Introduction

Cilia are microtubule-based organelles that project from the surface of most vertebrate cells and perform essential sensory and mechanical functions. They are built on a conserved microtubule scaffold, the axoneme, which is enclosed by the ciliary membrane and anchored to a centriole-derived basal body [1, 2]. Primary cilia are typically non-motile, display a 9+0 arrangement of microtubule doublets, and act as signaling hubs for extracellular cues [3]. In contrast, motile cilia exhibit a 9+2 architecture and use axonemal motility machinery to drive fluid and particle movement across epithelial surfaces [4–6]. Despite these structural and functional specializations, primary and motile cilia rely on structural and regulatory modules that assemble and maintain a functional axoneme.

The physiological importance of cilia is underscored by ciliopathies, a broad spectrum of multisystem genetic disorders resulting from defects in ciliary assembly, structure, or function [7–9]. Traditionally, ciliopathies are divided into sensory ciliopathies, such as Joubert syndrome, which present with neurodevelopmental, retinal, and renal phenotypes, and motile ciliopathies, such as primary ciliary dyskinesia (PCD), which are characterized by respiratory and laterality defects. Emerging genetic and clinical data increasingly challenge this classification by revealing overlap between these categories, both within individuals and at the gene level [10–15]. In some cases, pathogenic variants in a single gene cause both sensory and motile cilia phenotypes within the same individuals. For example, variants in KIAA0586/TALPID3 have been associated with Joubert syndrome features and chronic airway disease, whereas variants in IFT74 have been shown to cause a PCD-like respiratory phenotype together with retinal and skeletal abnormalities [12, 16]. More commonly, overlap is supported at the gene level, as factors classically linked to primary ciliopathies have also been implicated in motile cilia phenotypes in separate cohorts or functional studies. Examples include OFD1 [17–19], NEK10 [20, 21], CCDC57 [22–24], IFT140 [25] and CEP164 [26]. Collectively, these findings suggest that ciliogenesis relies on shared molecular modules, including microtubule-associated proteins, that govern organelle biogenesis and function across both primary and motile cilia.

A mechanistic understanding of this clinical overlap requires defining how proteins implicated in sensory ciliopathies contribute to motile ciliogenesis in multiciliated cells (MCCs). Vertebrate multiciliated epithelia generate motile cilia arrays adapted to tissue-specific needs [27]. For example, *Xenopus laevis* embryonic epidermal MCCs are interspersed within the epidermis and generate directed cilia-driven fluid flow across the embryo surface [28–30]. In the airway, MCCs assemble dense ciliary arrays across a broad apical surface to drive mucociliary clearance, whereas ependymal MCCs lining the brain ventricles generate fewer cilia on smaller apical domains to propel cerebrospinal fluid [31, 32]. Despite these differences in cell geometry, cilia number, and array architecture, the core multiciliogenesis program is broadly conserved across vertebrate tissues [31–33]. Comparing MCCs across complementary model systems therefore provides a powerful approach to identify conserved regulators of multiciliogenesis and to understand why some genes linked to primary ciliopathies also give rise to motile cilia phenotypes.

MCC differentiation requires expansion of the basal body pool to build tens to hundreds of motile cilia [31–33]. This is achieved through two coordinated centriole amplification pathways (Fig. 1A). In the centriolar pathway, parental centrioles template new procentrioles. In parallel, the deuterosome-mediated *de novo* pathway produces the majority of centrioles through DEUP1-positive scaffolds that nucleate multiple procentrioles [34–38]. An MCC-specific assembly implicated in this process is PCM1-positive fibrogranular material (FGM), which associates with deuterosomes and influences their size, number, and distribution [37, 39, 40]. FGMs also concentrate centriole-associated proteins, including factors required for amplification, thereby promoting their proper targeting and supporting ciliary structure and motility [40].

**Figure 1:**
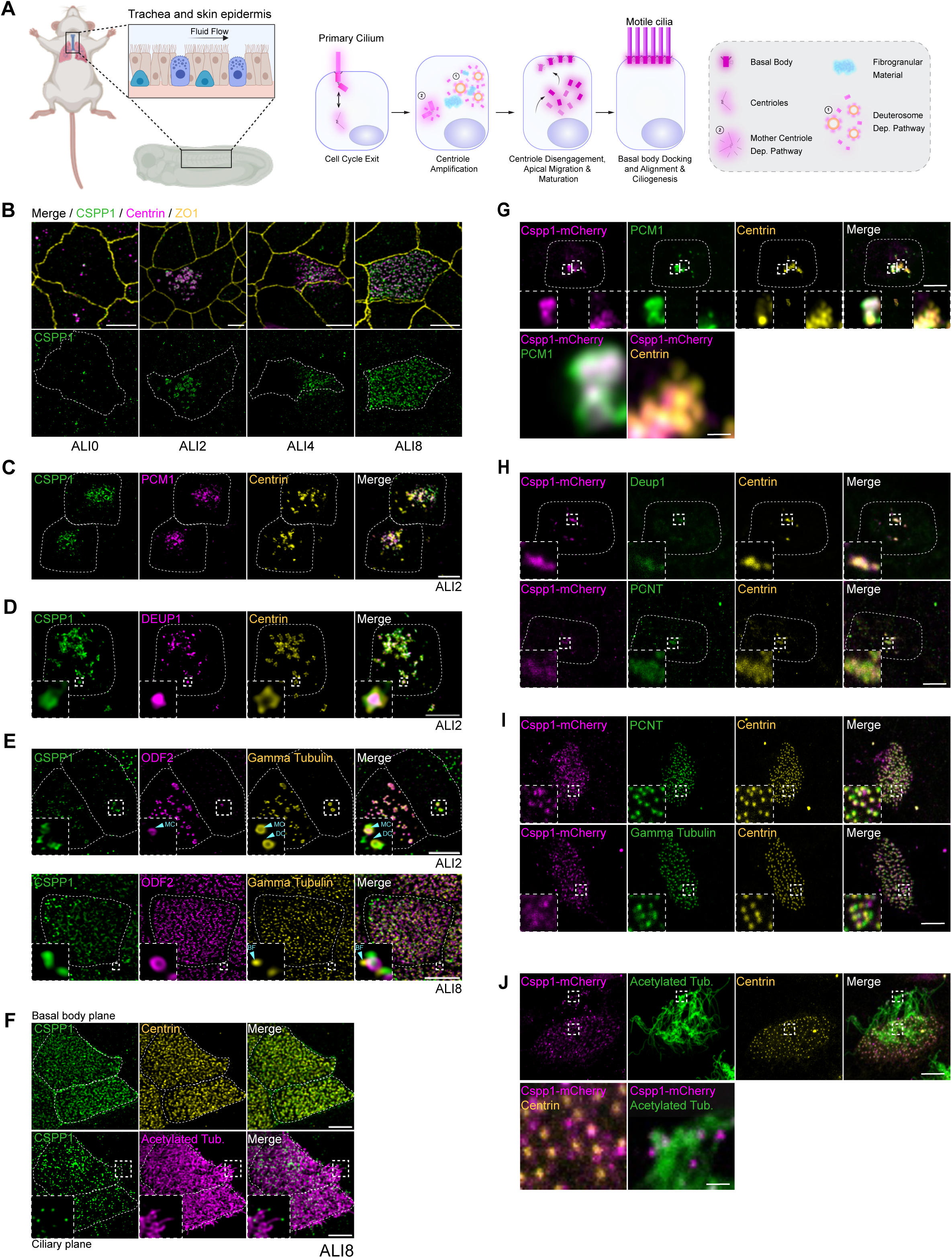
CSPP1 localizes to fibrous granules, deuterosomes, basal bodies and ciliary axonemes during multiciliogenesis. **(A)** Schematic illustration of multiciliated cell (MCC) differentiation in mouse tracheal epithelial cells (MTECs) and *Xenopus* embryonic epidermis. Key stages include cell cycle exit, centriole amplification (fibrous granules and deuterosomes), basal body docking, and ciliary axoneme elongation. (**B–F) CSPP1 localization in Mouse Tracheal Epithelial Cells (MTECs).** Representative immunofluorescence images of MTECs at different air-liquid interface (ALI) culture days. **Scale bars:** 5 μm. **(B) Centrioles.** Cells were stained for CSPP1 (green), Centrin (magenta; centrioles), and ZO-1 (yellow; tight junctions) across differentiation stages. **(C) Fibrous granules.** MTECs at ALI2 (amplification) stained for CSPP1 (green), PCM1 (magenta; fibrous granules), and Centrin (yellow). **(D) Deuterosomes.** At ALI2, CSPP1 (green) associates with deuterosome cores marked by DEUP1 (magenta), surrounded by Centrin-positive nascent centrioles (yellow). Inset in ALI2 is 2.5X zoom. **(E) Proximal-distal orientation at the centrioles.** Top (ALI2) and Bottom (ALI8): CSPP1 (green) localizes relative to ODF2 (magenta; mother centriole appendage) and gamma-tubulin (yellow). Cyan arrowheads indicate the mother centriole (MC); DC indicates daughter centriole. Inset in ciliary plane is 2.5X zoom. **(F) Basal bodies and cilia (ALI8).** Orthogonal views show CSPP1 (green) localizing to the basal body plane (co-stained with Centrin, yellow) and extending into the ciliary plane (co-stained with Acetylated Tubulin, magenta). Inset in ciliary plane is 3X zoom. **(G–J) Cspp1–mCherry localization in *Xenopus* epidermal MCCs.** Representative immunofluorescence images of Xenopus MCCs at different differentiation stages. **Scale bars:** 10 μm (main panels), 0.1 μm (insets). **(G) Fibrous granules.** During early amplification, Cspp1–mCherry (magenta) co-localizes with PCM1 (green) and Centrin (yellow) within fibrous granules. Intercalating cell is outlined by dashed lines. Insets are 10X zoom. **(H) Deuterosomes.** During active amplification, Cspp1–mCherry (magenta) associates with deuterosomes marked by Deup1 (top, green) and PCNT (bottom, green), surrounded by Centrin-positive nascent centrioles (yellow). Intercalating cells are outlined by dashed lines. Inset is 10X zoom. **(I) Basal bodies.** In differentiated MCCs, Cspp1–mCherry (magenta) localizes to basal bodies marked by PCNT (top, green) and gamma-tubulin (bottom, green), occupying a region partially distinct from the Centrin core (yellow). Inset is 10X zoom. **(J) Cilia.** Cspp1–mCherry (magenta) localizes to the ciliary axoneme marked by Acetylated Tubulin (green) with enrichment at the distal tips, while maintaining basal body localization (bottom panels; Centrin in yellow).

Following amplification, nascent centrioles elongate and mature, disengage from deuterosomes and parental centrioles, migrate toward the apical cortex, and dock at the plasma membrane to act as basal bodies that template axoneme assembly. These transitions require coordinated remodeling of the actin and microtubule cytoskeletons. Actin networks contribute to basal body docking and apical spacing, while apical microtubules support coordinated planar alignment downstream of planar cell polarity cues [41–44]. Recent imaging in brain MCC progenitors further supports a role for microtubules in organizing centriole amplification intermediates and promoting the collective repositioning and apical migration of newly formed centrioles [45]. Together, these observations suggest that microtubule-associated proteins are well positioned to couple centriole amplification to later steps of basal body organization and axoneme assembly.

In this context, CSPP1 emerges as a strong candidate and a potential shared regulator of primary and motile ciliogenesis. *In vitro*, CSPP1 acts similarly to microtubule-stabilizing agents by binding to growing microtubule ends and attenuating their dynamics [46, 47]. In mammalian cells, CSPP1 promotes axoneme elongation and the formation of Hedgehog signaling–competent cilia, and it has also been implicated in cell division and adhesion [48–50]. Clinically, pathogenic variants in CSPP1 cause the neurodevelopmental disorder Joubert syndrome and can also underlie the more severe Meckel–Gruber syndrome [51–53]. Notably, a subset of individuals with Joubert syndrome harboring CSPP1 variants present with respiratory manifestations such as chronic sinusitis [53]. CSPP1 links to motile cilia are further supported by the hydrocephalus phenotype observed upon *cspp1* loss in zebrafish and by comparative proteomics identifying CSPP1 as a conserved component of the motile ciliary proteome across diverse species [48, 54]. Despite these associations, the roles of CSPP1 in centriole amplification, basal body organization, and axoneme assembly during multiciliogenesis remain unknown.

Here, we use mouse tracheal epithelial cell cultures and *Xenopus laevis* embryonic epidermis as complementary MCC models to elucidate CSPP1 function during multiciliogenesis. We show that CSPP1 localizes to fibrous granules and deuterosomes during centriole amplification and to basal bodies and the distal axoneme in mature MCCs. Using loss-of-function approaches, we find that CSPP1 is required for efficient centriole amplification, basal body apical docking and organization, and the formation of structurally intact and functional motile cilia. These findings establish CSPP1 as a conserved component of the vertebrate multiciliogenesis machinery.

## Results

### CSPP1 is a conserved component of the multiciliogenesis machinery in vertebrate MCCs

To test whether CSPP1 is a conserved component of the multiciliogenesis machinery, we mapped its localization across multiciliated cell models and tissues. We focused on mouse tracheal epithelial cells (MTECs) undergoing synchronized differentiation *in vitro* and on Xenopus embryonic epidermis *in vivo*, and extended these analyses to mouse trachea, ependyma, and human airway epithelia. Comparative proteomic analyses previously identified CSPP1 as an ancestral ciliary protein in organisms ranging from choanoflagellates to sea anemones and sea urchins [54]. Multiple sequence alignment further showed that CSPP1 is highly conserved across vertebrates (Fig. S1A). CSPP1 also contains helical regions distributed throughout the protein, central microtubule-binding and pausing domains implicated in microtubule stabilization, and a C-terminal centrosome-targeting domain (Fig. S1A) [46, 48]. We therefore examined CSPP1 localization relative to centrioles and basal bodies, centriole amplification structures, and cilia during multiciliogenesis (Fig. 1A).

In the MTEC culture system, the transition to an air–liquid interface (ALI) triggers a staged differentiation program (Fig. 1A). Following cell-cycle exit, early differentiation (ALI1–2) is characterized by formation of fibrous granules and deuterosomes and initiation of centriole amplification. Centrioles then elongate (ALI3–4), disengage and migrate apically (ALI4–8), and finally dock at the plasma membrane to template axoneme assembly (ALI8–12) (Fig. 1A). To define CSPP1 localization across these stages, we performed 3D structured illumination microscopy (3D-SIM) and co-stained cells with markers for centrioles and basal bodies (Centrin, ODF2), fibrous granules (PCM1), deuterosomes (DEUP1 and PCNT), and cilia (acetylated tubulin and ARL13B), using the tight junction marker ZO-1 to delineate apical boundaries. Gamma-tubulin was used to mark the centrosome during early stages and the basal foot after basal body docking.

CSPP1 localized to centrioles throughout MTEC differentiation (Fig. 1B). At ALI2, CSPP1 localized to cytoplasmic puncta associated with PCM1- and DEUP1-positive structures, showing partial overlap with fibrous granules and deuterosomes (Fig. 1C, 1D). Notably, CSPP1 formed a ring-like structure surrounding the DEUP1 core, a pattern recapitulated in cells expressing mNG-CSPP1 (Fig. S1C). We next resolved CSPP1 relative to centriolar markers at ALI2 and ALI8. At ALI2, CSPP1 occupied a domain closely associated with, but distinct from, the gamma-tubulin-positive centrosome and ODF2-positive subdistal appendages. After apical docking (ALI8), CSPP1 localized to a domain distal to ODF2 and distinct from the gamma-tubulin-positive basal foot (Fig. 1E). In fully ciliated cells, CSPP1 remained associated with basal bodies and was enriched at the distal region of the ciliary axoneme (Fig. 1F).

To validate these *in vitro* observations in tissue, we next examined *ex vivo* mouse trachea and ependyma. In the tracheal lumen, CSPP1 localized to basal bodies and cilia, and detached cilia revealed an axonemal pool with prominent distal tip enrichment (Fig. S1B). A comparable distribution was observed in mouse ependymal cells (Fig. S1D). During amplification, CSPP1 associated with gamma-tubulin- and PCNT-positive deuterosomes and with Centrin-positive nascent centrioles. In mature ependymal MCCs, CSPP1 localized to basal bodies and the distal region of the axoneme. To extend these observations to human cells, we analyzed human airway epithelial cultures, where CSPP1 similarly localized to basal bodies and the ciliary distal tip (Fig. S1E).

We next examined Cspp1 localization across distinct stages of multiciliogenesis in the *Xenopus* embryonic epidermis expressing Cspp1–mCherry (Fig. 1G–J). Using a newly generated and validated *Xenopus*-specific anti-PCM1 antibody (Fig. S1F), we found that Cspp1 co-localized with PCM1 at fibrous granules during early differentiation. Co-staining with Centrin revealed two fibrous granule populations: a Centrin-negative pool, consistent with early fibrous granules before centriole amplification, and a Centrin-positive pool, indicating that Cspp1 is retained at fibrous granules engaged in procentriole assembly (Fig. 1G). Because these active fibrous granules are coupled to the deuterosome pathway, we next assessed Cspp1 localization relative to deuterosomes and found that Cspp1-mCherry strongly co-localized with Deup1- and PCNT-positive deuterosomes surrounded by nascent centrioles (Fig. 1H). In mature ciliated MCCs, apical imaging showed that Cspp1-mCherry localized to the basal body region, where it occupied a domain closely associated with, but partially distinct from, Centrin, PCNT, and gamma-tubulin (Fig. 1I). Imaging in the ciliary plane further revealed prominent enrichment of Cspp1 at the distal tips of ciliary axonemes (Fig. 1J).

Together, these findings establish CSPP1 as a conserved component of the multiciliogenesis machinery in mouse, human and *Xenopus*. Its localization to fibrous granules and deuterosomes during centriole amplification, and to basal bodies and ciliary axonemes in mature cells, suggests potential roles in centriole amplification and later steps of motile cilia assembly and function.

### CSPP1 is required for efficient centriole amplification in mouse and *Xenopus* MCCs

We next tested whether CSPP1 is required for centriole amplification using loss-of-function approaches in both MTECs and *Xenopus* epidermal multiciliated cells.In MTECs, we depleted CSPP1 using lentiviral shRNA-mediated knockdown. Knockdown efficiency was first validated by Western blotting, which showed a strong reduction in CSPP1 protein levels (Fig. S2A). CSPP1 depletion was then confirmed by immunofluorescence in differentiating MTECs following lentiviral infection and puromycin selection (Fig. S2B). To assess centriole amplification, we stained control and shCspp1-transduced cultures for Centrin and ZO-1 at multiple differentiation stages.

CSPP1 depletion impaired centriole amplification across complementary readouts of frequency and morphology. At ALI4, the fraction of cells that underwent centriole amplification, defined by the presence of multiple Centrin-positive foci, was reduced by 27.6% compared with controls (Fig. 2A, 2B). This reduction persisted at ALI8, indicating impaired centriole amplification rather than a delay in differentiation (Fig. 2A, 2B). Among the subset of shCspp1 cells that underwent centriole amplification, the resulting apical centriole arrays were also frequently abnormal. Rather than forming the dense, uniform distribution seen in controls, these cells exhibited reduced centriole density, disorganized spatial organization, and elongated Centrin filaments (Fig. 2A). Quantification confirmed that these defects were substantially more frequent upon CSPP1 depletion (Fig. 2C). Together, these abnormal features may reflect defects in centriole amplification as well as impaired progression into later stages of MCC differentiation. We addressed these possibilities in subsequent analyses.

**Figure 2.**
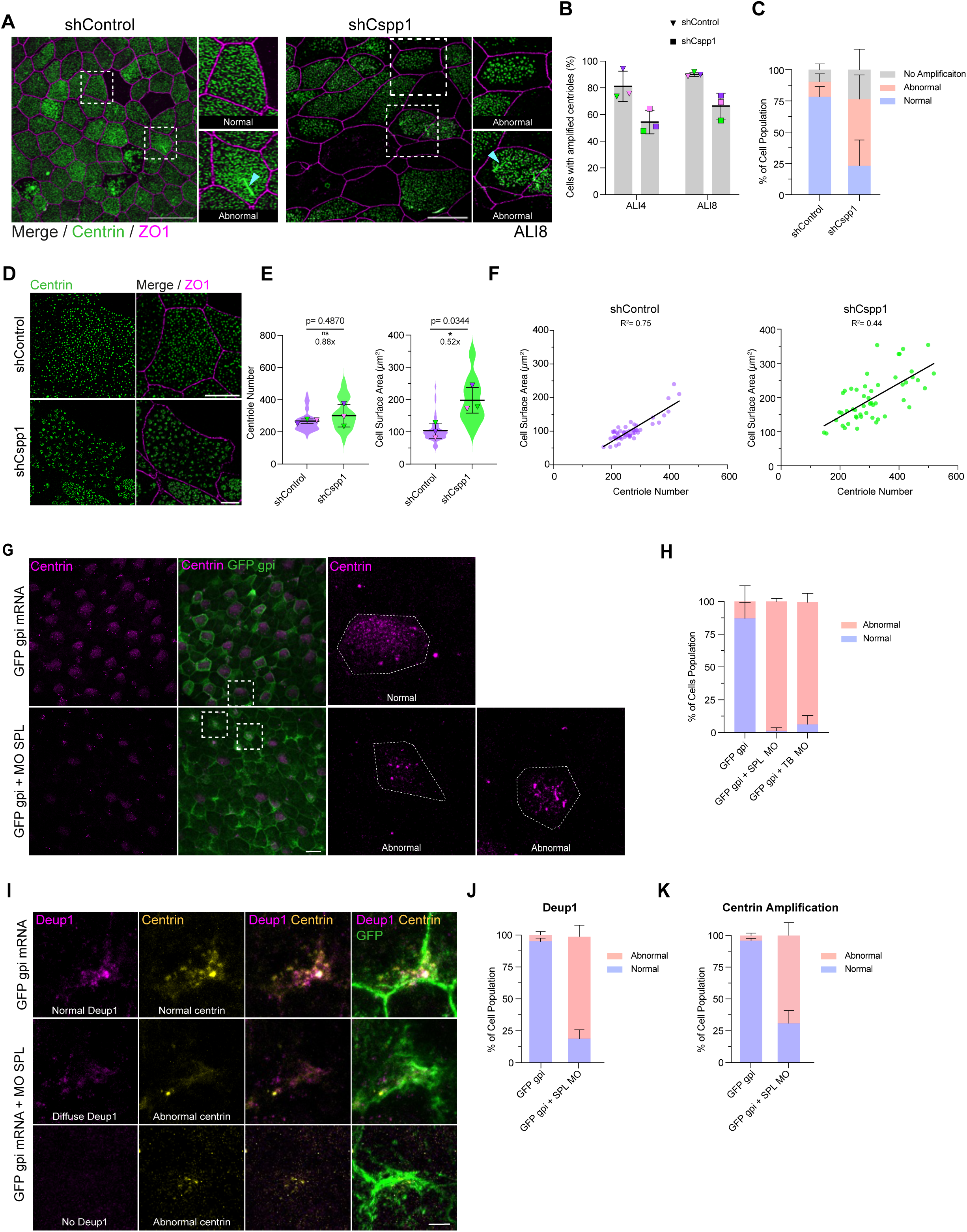
CSPP1 is required for efficient centriole amplification and deuterosome assembly. **(A) Centriole amplification phenotypes in MTECs.** Representative immunofluorescence images of ALI8 MTEC cells transduced with Control (shScramble) or *Cspp1* (shCspp1) shRNA lentivirus. Cells were stained for Centrin (green) and ZO-1 (magenta). Control cells display a **uniform apical distribution** of basal bodies. *Cspp1*-depleted cells exhibit abnormal phenotypes, characterized by sparse, irregular distributions or Centrin **aggregates and filaments** (cyan arrow). Scale bar: 20 μm. **(B) Quantification of centriole amplification efficiency.** The percentage of cells with amplified centrioles was quantified at early (ALI4) and late (ALI8) stages. *Cspp1* depletion significantly impairs amplification. Data represent mean ± SD from three biological replicates (n=30 fields total; 10 fields per replicate, containing ∼50–120 cells per field). Statistical significance determined by paired, two-tailed Student’s t-test. **(C) Quantification of basal body amplification phenotypes.** Among cells that successfully generated centrioles (quantified in B), phenotypes were categorized as “Normal” (discrete, spread-out centrioles) or “Abnormal” (characterized by aggregates, filaments, or aberrant clustering). *Cspp1* depletion results in a significant increase in the proportion of cells displaying aberrant morphology. Data represent mean ± SD from three biological replicates (n=30 fields analyzed). Statistical significance determined by two-way ANOVA. **(D) High-resolution analysis of basal body organization.** Representative 3D-SIM images of ALI8 MTECs. Cells were stained for Centrin (green) and ZO-1 (magenta). High-resolution structured illumination microscopy (SIM) was used to resolve individual centrioles, enabling precise quantification of centriole amplification efficiency. Scale bar: 5 μm. **(E) Quantification of centriole number and apical surface area.** Violin plots display the distribution of centriole numbers (left) and apical surface area (right) per cell. Notably, while the total centriole number per cell remained unchanged (p=0.4870), *Cspp1*-depleted cells exhibited a significant increase in apical surface area (p=0.0344) compared to controls. Magenta, pink, and green triangles represent means of independent biological replicates (n=51 cells total). Statistical significance determined by paired, two-tailed Student’s t-test. *P<0.05, ns: not significant. **(F) Correlation of centriole amplification with apical cell surface area.** Centriole number was plotted against apical surface area for individual cells quantified in (D). Linear regression analysis reveals a strong positive correlation in control cells between centriole numbers and apical cell surface area(R² = 0.75). This scaling relationship is weakened in *Cspp1*-depleted cells (R² = 0.44). **(G) Centriole amplification defects in *Xenopus*.** Representative confocal images of *Xenopus* embryonic epidermis at stage 30 (st30), injected at the 4-cell stage with the the splice-blocking Cspp1 morpholino (MO SPL). Cells were stained for GFP (lineage tracer, green) and Centrin (magenta). Control MCCs show a **uniform apical distribution** of centrioles, whereas *Cspp1* morphants display irregular distribution and distinct Centrin-marked **aggregates and filaments**. Scale bar: 20 μm. Inset is 5X zoom. The apical cell surface is outlined with dashed lines. **(H) Quantification of basal body distribution phenotypes.** “Abnormal” is defined by non-uniform distribution or the presence of aggregates. Data include results from both splice-blocking (MO SPL) and translation-blocking (MO TB) morpholinos. Data represent mean ± SD from three biological replicates (n>1000 cells analyzed from >14 embryos per condition). Statistical significance determined by one-way ANOVA (Dunnett’s test).. **(I) Deuterosome formation in *Xenopus* MCCs.** Representative images of deuterosome formation in *Xenopus* MCCs at stage 17 (st17). Cells were stained for Deup1 (magenta), Centrin (yellow), and GFP (green). Control cells show robust Deup1-positive deuterosomes surrounded by nascent centrioles. *Cspp1* morphants display diffuse or absent Deup1 signal and disorganized Centrin puncta (“Abnormal”). Scale bar: 5 μm. **(J) Quantification of Deup1 phenotypes.** Deuterosome formation was categorized as “Normal” (discrete puncta) or “Abnormal” (diffuse/absent). Data represent mean ± SD from three biological replicates (n>2000 cells analyzed from >11 embryos per condition). Statistical significance determined by unpaired Welch’s t-test. **(K) Quantification of Centrin amplification defects.** Centrin signal was categorized as “**Normal**” (discrete puncta around deuterosomes), “**Abnormal**”, or **“No Amplification”** (diffuse/absent). Data represent mean ± SD from three biological replicates (n>2000 cells analyzed from >11 embryos per condition). Statistical significance determined by unpaired Welch’s t-test. *P<0.05; **P<0.01, ***P<0.001, ****P<0.0001; ns, not significant.

To determine whether the reduced centriole density reflected altered centriole number, we quantified centriole number at ALI8, when centrioles are fully disengaged and can be resolved. Mean centriole number per cell was comparable between control and shCspp1 cells, but apical area was significantly increased after CSPP1 depletion (Fig. 2D, 2E). Consistent with the scaling relationship reported for airway and *Xenopus* MCCs [55, 56], centriole number correlated strongly with apical area in control cells (R² = 0.75), but this relationship was weaker in shCspp1 cells (R² = 0.44) (Fig. 2F). Thus, the reduced centriole density is driven by the increased apical area without a compensatory increase in centriole amplification. Together with the decreased amplification frequency (Fig. 2B), these data indicate that CSPP1 promotes efficient centriole amplification and proper scaling of centriole number to apical domain size.

To test whether this requirement is conserved *in vivo*, we depleted Cspp1 in *Xenopus* embryos by injecting the presumptive epidermis with translation-blocking (TB) or splice-blocking (SPL) morpholinos (Fig. S2C). Knockdown efficiency was validated by RT-PCR for the SPL morpholino (Fig. S2D) and by silencing of a Cspp1–mCherry reporter for the TB morpholino (Fig. S2E). Because 30% of Cspp1 morphant MCCs failed to intercalate into the outer epidermal layer, we restricted all subsequent analyses to successfully intercalated cells (Fig. 2G, S2E). This ensured that the phenotypes described below reflect CSPP1 depletion rather than secondary consequences of impaired intercalation.

At developmental stage 30, control MCCs displayed a dense and uniform apical centriole distribution. In contrast, Cspp1 morphants exhibited severe centriole amplification defects. Many morphant cells failed to amplify centrioles. Consistent with our observations in MTECs, among the cells that did undergo amplification, the resulting centriole arrays were frequently abnormal, characterized by disorganized Centrin aggregates, elongated Centrin-positive filaments, or sparse and irregular centriole distributions across the apical domain. These defects were consistently observed with both the SPL morpholino (Fig. 2G) and the TB morpholino (Fig. S2F). Quantification confirmed that nearly all intercalated morphant cells fell into these abnormal categories (Fig. 2H).

To define where Cspp1 acts within the amplification pathway, we examined the deuterosome-dependent program, which generates the majority of centrioles in MCCs [57]. At developmental stage 17, the peak of centriole amplification, control cells contained robust DEUP1-positive deuterosomes surrounded by nascent centrioles. In contrast, Cspp1 morphants showed a striking loss of deuterosomes, with DEUP1 signal appearing diffuse or absent, coincident with defective centriole amplification (Fig. 2I–K). To determine whether this defect arose further upstream, we next assessed fibrous granules [37, 38, 58, 59]. Using a *Xenopus*-specific anti-PCM1 antibody, we observed discrete fibrous granules in both control and Cspp1 morphant cells. Quantification of PCM1 granule number and size revealed no significant differences between conditions (Fig. S2G, S2H). Together, these data indicate that Cspp1 is dispensable for initial fibrous granule assembly but is strictly required downstream for proper deuterosome formation.

Collectively, these data from mouse and *Xenopus* demonstrate a conserved requirement for CSPP1 in centriole amplification. Across both systems, CSPP1 depletion reduced the efficiency of centriole amplification and disrupted formation of the dense centriole arrays characteristic of MCCs.

### CSPP1 is required for basal body apical migration, spatial organization and rotational polarity

Having established that CSPP1 is required for centriole amplification, we next asked whether it also contributes to later steps of multiciliogenesis, specifically basal body apical migration, spatial organization, and rotational polarity.

To assess apical migration, we evaluated the depth of basal bodies relative to the apical membrane marker ZO-1 using 3D reconstructions and orthogonal (X-Z) projections at ALI20 (Fig. 3A). In control cells, the majority of basal bodies reached the apical surface. In contrast, many basal bodies in shCspp1 cells remained stalled deeper within the cytoplasm (Fig. 3A). Quantification of the vertical distance between basal bodies and ZO-1 confirmed a significant apical migration defect, with mean basal body depth increasing from ∼1.2 μm in controls to ∼1.8 μm after CSPP1 depletion (Fig. 3A).

**Figure 3.**
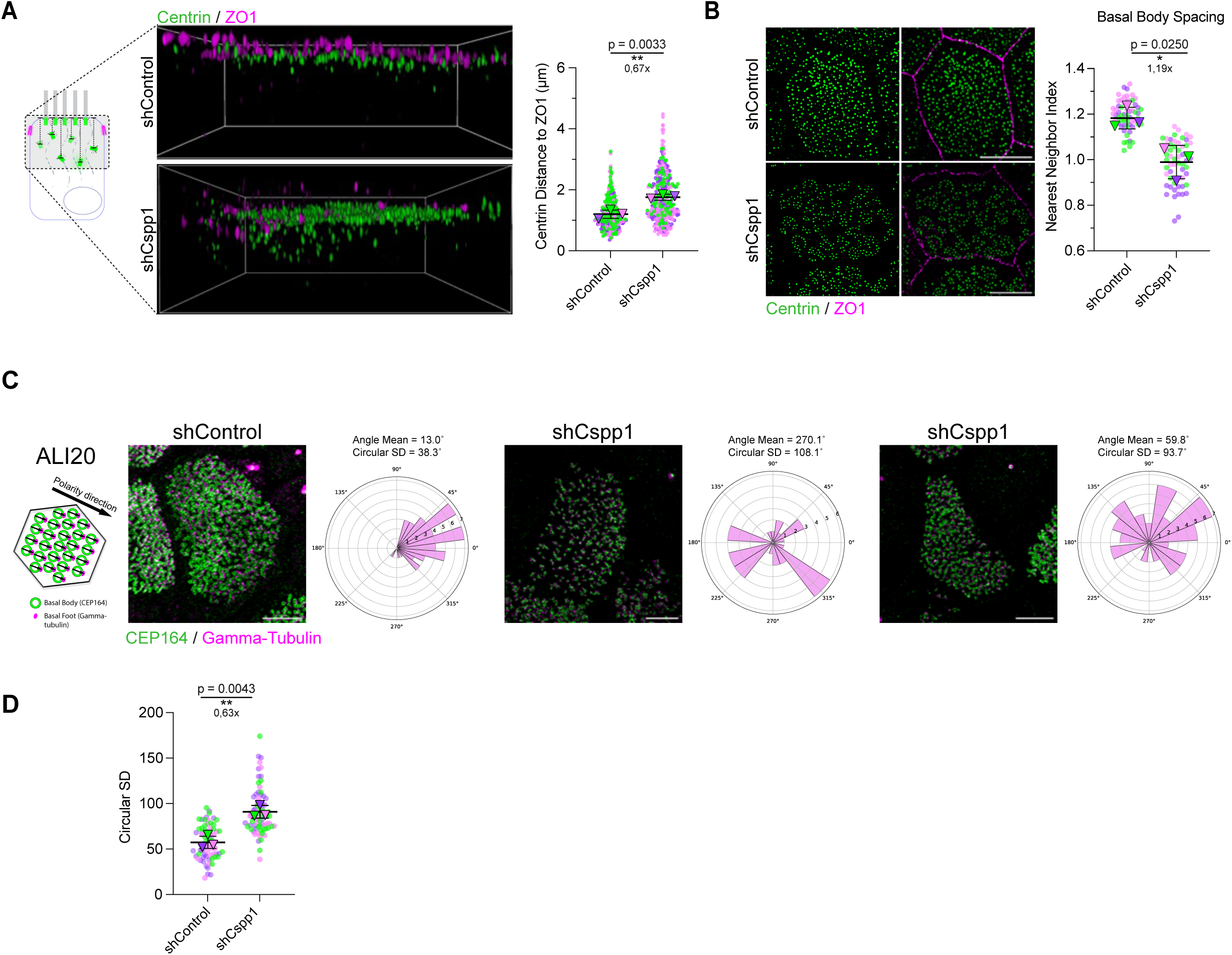
CSPP1 regulates basal body docking, organization, and rotational polarity. **(A) Basal body docking defects.** (Left) Representative 3D-SIM side-view reconstructions of ALI8 MTECs stained for Centrin (green) and ZO-1 (magenta). Images were processed with Aivia software to visualize basal body depth. (Right) Quantification of the vertical distance between basal bodies and the apical surface (ZO-1 plane). Measurements were restricted to basal bodies not colocalized with the ZO-1 plane. *Cspp1* depletion significantly impairs docking. Data represent mean ± SD from three independent biological replicates (n=30 cells analyzed; ∼300 basal bodies total). Statistical significance determined by unpaired Student’s t-test. **P=0.0033. **(B) Basal body apical organization.** (Left) Representative 3D-SIM images showing the spatial distribution of basal bodies at ALI8. (Right) Quantification of the Nearest Neighbor Index (NNI). Control cells exhibit a regular, dispersed distribution (higher NNI), whereas *Cspp1*-depleted cells show significant clustering (lower NNI). Data represent mean ± SD from three biological replicates (n=60 cells total). Statistical significance determined by paired Student’s t-test. *P=0.0250. Scale bar: 5 μm. **(C) Establishment of rotational polarity.** (Left) Schematic illustrating the determination of planar polarity vectors based on the angle between the basal body center (CEP164, green) and the basal foot (gamma-tubulin, magenta). (Right) Representative rose plots displaying the angular distribution of basal body vectors within individual ALI20 cells. Control cells show a uniform directionality (tight angular spread), while *Cspp1*-depleted cells exhibit disorganized polarity (broad angular spread). Scale bar: 2 μm. **(D) Quantification of rotational polarity.** The Circular Standard Deviation (CSD) was calculated for each cell to assess the consistency of basal body alignment. *Cspp1* depletion results in a significantly higher CSD, indicating disrupted rotational polarity. Data represent mean ± SD from three biological replicates (n=74 cells total). Statistical significance determined by paired Student’s t-test. **P=0.0043. *P<0.05; **P<0.01, ***P<0.001****, P<0.0001; ns, not significant.

We next asked whether the basal bodies that reached the apical domain were properly organized across the cell surface. In control cells, basal bodies formed a uniform apical array. In contrast, this organization was disrupted in shCspp1 cells that amplified centrioles, where basal bodies formed irregular clusters and left large regions of the apical surface unoccupied (Fig. 3B). We quantified this defect using the nearest-neighbor index (NNI), a measure of spatial regularity. Control cells exhibited a higher NNI (∼1.2), consistent with a more even pattern, whereas shCspp1 cells showed a significantly lower NNI (∼1.0), indicative of clustering and loss of regular spacing (Fig. 3B).

We then examined whether CSPP1 depletion affects basal body rotational polarity, which is necessary for coordinated effective stroke direction. At ALI20, we quantified basal body orientation using CEP164 to mark distal appendages and gamma-tubulin to mark the basal foot, defining a polarity vector from their relative positions for each basal body (Fig. 3C). In control cells, polarity vectors were tightly aligned, producing a narrow angular distribution. In contrast, shCspp1 cells showed a much broader angular spread and reduced within-cell alignment, resulting in a significantly higher circular standard deviation (mean SD ∼90.7°) compared to controls (∼57.4°) (Fig. 3D).

Together, these data show that CSPP1 is required not only for centriole amplification but also for subsequent basal body apical migration, spatial organization, and establishment of rotational polarity, all of which are important for assembly of a functional polarized ciliary array.

### CSPP1 depletion impairs cilia formation, length, and motility in MTECs

We next asked whether the defects in centriole amplification and basal body organization translate into defects in ciliogenesis. CSPP1 depletion significantly reduced the fraction of multiciliated cells at ALI8 (shControl ∼87.9% versus shCspp1 ∼68.9%; Fig. 4A, 4B). This reduction persisted at ALI12, indicating a sustained ciliogenesis defect rather than a simple delay in differentiation (Fig. S3A). Classification of ciliary phenotypes at ALI8 further revealed an increase in cells with abnormal or absent ciliation upon CSPP1 depletion (Fig. 4A, 4C). We therefore next examined the nature of these ciliary defects in more detail.

**Figure 4.**
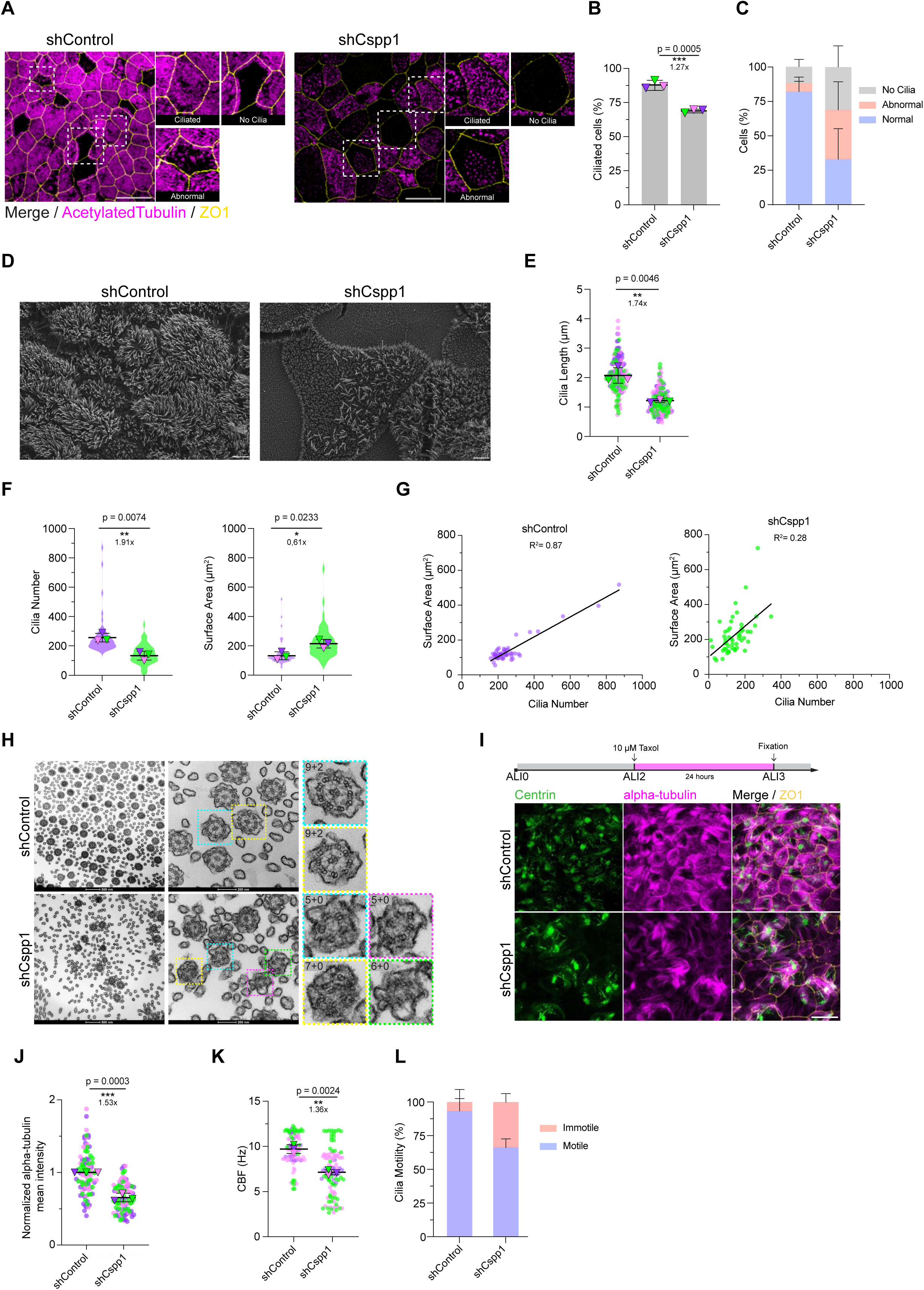
CSPP1 loss leads to defective cilia formation, axonemal structure, and motility. **(A) Cilia biogenesis defects.** Representative immunofluorescence images of ALI8 MTECs transduced with Control (shControl) or *Cspp1* (shCspp1) shRNA lentivirus. Cells were stained for Acetylated Tubulin (magenta) and ZO-1 (yellow). Control cells display dense, uniform ciliation. *Cspp1*-depleted cells exhibit abnormal phenotypes, characterized by sparse ciliation or a complete lack of cilia (“No Cilia”). Scale bar: 20 μm. **(B) Quantification of ciliation efficiency.** The percentage of ciliated cells at ALI8 was quantified. *Cspp1* depletion significantly reduces the proportion of ciliated cells. Data represent mean ± SD from three biological replicates (n=30 fields analyzed; 10 fields per replicate, containing ∼50–120 cells per field). Statistical significance determined by unpaired Student’s t-test. ***P=0.0005. **(C) Quantification of ciliary phenotypes.** Ciliated cell populations from (A) were categorized as “Normal” (homogeneous, full-length cilia) or “Abnormal” (sparse, short cilia). *Cspp1* depletion shifts the population significantly towards abnormal phenotypes. Data represent mean ± SD from three biological replicates (n=30 fields analyzed; 10 fields per replicate). Statistical significance determined by two-way ANOVA. ****P<0.0001, **P=0.0040. **(D) Scanning electron microscopy (SEM) analysis.** Representative SEM images of ALI8 MTECs used for high-resolution quantification of individual cilia. Control cells show dense carpets of cilia, whereas *Cspp1*-depleted cells display sparse, disorganized, and shortened cilia. Scale bar: 2 μm. **(E) Quantification of cilia length.** Cilia length was measured from SEM images. *Cspp1*-depleted cilia are significantly shorter than controls. Data represent mean ± SD from three biological replicates (n=51 cells total). Statistical significance determined by paired Student’s t-test. **P=0.0046. **(F) Quantification of cilia number and apical surface area.** Violin plots displaying the distribution of cilia number (left) and apical surface area (right) per cell based on SEM analysis. *Cspp1* depletion significantly reduces cilia number (**P=0.0074) despite a concurrent increase in apical surface area (*P=0.0233). Data represent mean ± SD from three biological replicates (n=50 cells total). Statistical significance determined by paired Student’s t-test. **(G) Correlation of cilia number with surface area.** Linear regression analysis of cilia number plotted against cell surface area. The strong positive correlation observed in control cells (R² = 0.87) is disrupted in *Cspp1*-depleted cells (R² = 0.28), indicating disruption in scaling cilia number with apical cell surface area. Data derived from the n=50 cells quantified in (E). **(H) Ultrastructural analysis of axonemes.** Transmission electron microscopy (TEM) images of transverse cilia sections. (Left) Low-magnification views of the ciliary field. (Right) High-magnification views of individual axonemes. Control cilia exhibit the canonical 9+2 microtubule arrangement. *Cspp1*-depleted cilia display severe structural defects, including loss of central pairs (9+0) and aberrant microtubule numbers (5+0, 6+0, 7+0). Scale bars: 500 nm (left), 200 nm (right). **(I) Apical microtubule network organization.** (Top) Schematic of the Taxol stabilization assay. MTECs were treated with Taxol (10 μM) from ALI2 to ALI3. (Bottom) Representative immunofluorescence images stained for Centrin (green), alpha-tubulin (magenta), and ZO-1 (yellow). *Cspp1*-depleted cells display a disorganized and less dense apical microtubule network. Scale bar: 20 μm. **(J) Quantification of apical microtubule intensity.** Mean fluorescence intensity of the Taxol-stabilized apical microtubule network was quantified from (I). Data represent mean ± SD from three biological replicates (n=30 image fields total; 10 fields per replicate). Statistical significance determined by paired Student’s t-test. ***P=0.0003. **(K) Cilia Beat Frequency (CBF).** CBF was measured in Hertz (Hz) using high-speed video microscopy at ALI8. *Cspp1*depletion results in a significant reduction in beat frequency. Data represent mean ± SD from three biological replicates (n=100 different fields analyzed). Statistical significance determined by paired Student’s t-test. **P=0.0024. **(L) Cilia Motility Analysis.** Quantification of the percentage of motile vs. immotile cilia within the population derived from the CBF analysis. *Cspp1* depletion significantly increases the fraction of immotile cilia. Data represent mean ± SD from three biological replicates (n=3). Statistical significance determined by paired Student’s t-test. **P=0.0069 (Motile), **P=0.0069 (Immotile). *P<0.05; **P<0.01, ***P<0.001, ****P<0.0001; ns, not significant.

Among the cells that did form cilia, shCspp1 cultures frequently displayed sparse, irregular, and shorter ciliary arrays, whereas control cultures showed dense and uniform ciliation (Fig. 4A). Scanning electron microscopy and subsequent quantification confirmed that shCspp1 cells extended significantly fewer and shorter cilia per cell than controls (Fig. 4D-F). Because mean centriole number at ALI8 was comparable between conditions in the subset of celsl that successfully amplified (Fig. 2D), these findings indicate that not all available centrioles efficiently generated cilia after CSPP1 depletion. Consistent with this, although shCspp1 cells exhibited enlarged apical surface areas (Fig. 4F), the relationship between apical area and cilia number was markedly weaker than in controls (control R² = 0.87; shCspp1 R² = 0.28; Fig. 4G). This disruption was stronger than that observed for the apical area-centriole number relationship (Fig. 2F). Together, these data indicate that CSPP1 is required for efficient cilia formation and normal axoneme elongation.

To determine whether these defects in cilia number and length were accompanied by abnormalities in axoneme structure, we analyzed ciliary ultrastructure by transmission electron microscopy (TEM) on transverse sections. Control MTECs consistently exhibited the canonical 9+2 organization of motile cilia, with nine outer doublets surrounding a central pair. In contrast, shCspp1 cilia displayed severe ultrastructural defects, frequently losing 9-fold symmetry, with examples including 5+0, 6+0, and 7+0 arrangements, and often lacking the central pair apparatus (Fig. 4H).

Given that CSPP1 is a microtubule-associated protein, we next asked whether the axonemal defects were accompanied by alterations in the apical microtubule network. To visualize this network, we stabilized microtubules with Taxol prior to fixation, a treatment that reveals the cortical meshwork typically obscured by ciliary tubulin [20]. In control cells, this exposed a robust and highly organized apical microtubule network. In contrast, shCspp1 cells displayed a sparser and less organized apical microtubule pattern, with reduced overall microtubule intensity and loss of the coherent directionality seen in controls (Fig. 4I, 4J). These findings indicate that CSPP1 depletion disrupts apical microtubule organization, consistent with a role for CSPP1 in the cytoskeletal architecture that supports efficient axoneme assembly and maintenance.

Finally, we examined the functional consequences of these structural and cytoskeletal defects by measuring ciliary motility. High-speed video microscopy of ALI cultures showed that ciliary beating was significantly impaired in shCspp1 cells (Movie S1) relative to control cells (Movie S2). Control cultures exhibited a mean ciliary beat frequency (CBF) of approximately 9.7 Hz, whereas shCspp1 cultures showed reduced motility with a mean CBF of around 7.1 Hz (Fig. 4K). Notably, a substantial fraction of cilia in depleted cultures beat below 5 Hz or appeared immotile (Fig. 4L). Together, these findings indicate that CSPP1 is required for the robust formation of structurally intact motile cilia and for maintaining normal ciliary beating.

### Cspp1 loss impairs ciiogenesis and cilia-driven flow *in vivo*

To evaluate ciliogenesis and cilia-driven transport *in vivo*, we next used the *Xenopus* embryonic epidermis. Consistent with the centriole amplification defects, Cspp1 depletion severely impaired cilia formation at stage 30 (Fig. 5A, Fig. S3B). While control MCCs displayed densely ciliated apical surfaces, successfully intercalated Cspp1 morphant cells frequently failed to generate normal ciliary arrays, forming either no cilia or only sparse, disorganized axonemes (Fig. 5A; Fig. S3B). Quantification confirmed that the vast majority of intercalated morphant cells fell into these abnormal categories (Fig. 5B).

**Figure 5.**
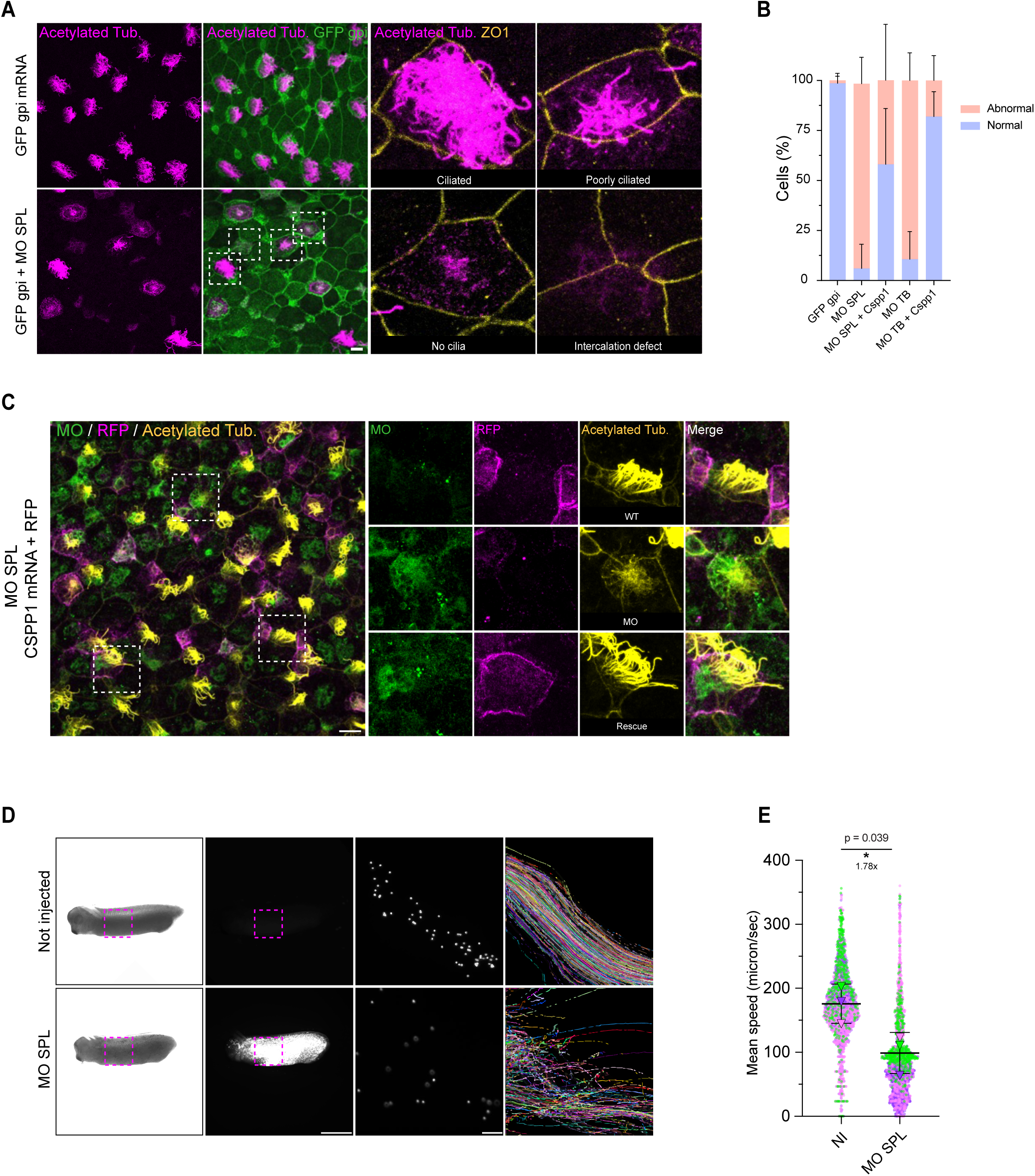
*Cspp1* loss leads to defective cilia formation and mucociliary transport in *Xenopus laevis*. **(A) Ciliogenesis profiles at Stage 30.** Representative confocal images of the *Xenopus* embryonic epidermis injected at the 4-cell stage with the indicated constructs. Cells were stained for the lineage tracer GFP (green), Acetylated alpha-tubulin (cilia, magenta), and ZO-1 (apical junctions, yellow). Control cells (GFP-GPI mRNA) display a uniform, dense ciliary array. *Cspp1* morphants (MO SPL) exhibit severe defects, classified as “Poorly ciliated,” “No cilia,” or “Intercalation defect.” Scale bar: 20 μm. Inset is a 5X zoom **(B) Quantification of ciliary phenotypes.** The distribution of ciliary phenotypes (“Normal” vs. “Abnormal”) was quantified. “Abnormal” includes cells with sparse, short, or absent cilia. The analysis includes both Splice-blocking (SPL) and Translation-blocking (TB) morpholinos, as well as rescue conditions. *Cspp1* depletion significantly increases the proportion of abnormal cells, which is rescued by co-injection of *Cspp1* mRNA. Data represent mean ± SD from three biological replicates (n > 500 cells analyzed from >13 embryos per condition). Statistical significance determined by One-way ANOVA (Šidák’s test). **(C) Rescue of ciliogenesis defects.** (Left) Low-magnification view of Stage 30 epidermis co-injected with *Cspp1* MO (SPL) and MO-resistant *Cspp1* mRNA + RFP tracer. (Right) High-magnification split-channel views. Cells were stained for the MO lineage tracer (green), the Rescue tracer (RFP, magenta), and cilia (Acetylated alpha-tubulin, yellow). Cells containing only the MO (green only) display the “Abnormal” phenotype, whereas cells successfully co-injected with the rescue construct (green + magenta) display “WT”-like ciliogenesis (“Rescue”). Scale bar: 20 μm. Inset (rigth-hand side) is a 3X zoom. **(D) Analysis of mucociliary flow.** Representative analysis of fluid flow across the embryonic epidermis. Fluorescent microbeads were applied to the surface of Stage 30 embryos. (Columns 1–2) Brightfield and fluorescent images of whole embryos showing the region of interest (ROI, yellow box). (Columns 3–4) Magnified views of the ROI showing bead positions and their cumulative trajectories (Kalman tracks) over time. Control (Not Injected, NI) embryos show long, directional flow trajectories. *Cspp1* morphants display short, disorganized tracks indicative of defective flow. Scale bars: 1 mm (embryo) and 50 μm (tracks). **(E) Quantification of flow speed.** Mean bead speed (µm/sec) was calculated from the tracking data in (D). *Cspp1* depletion significantly reduces the speed of mucociliary flow. Data represent mean ± SD from three biological replicates (n > 2100 beads measured from >8 embryos per condition). Statistical significance determined by Kruskal-Wallis test. *P=0.039. *P<0.05; **P<0.01, ***P<0.001, ****P<0.0001; ns, not significant.

To confirm specificity, we performed rescue experiments using MO-resistant RFP-Cspp1 mRNA. We first titrated the mRNA to identify a dose that did not cause toxicity or ciliary defects in wild-type embryos (Fig. S3C). Co-injection of Cspp1 mRNA with either the SPL or TB morpholino restored dense apical ciliogenesis and reduced the proportion of abnormal cells (Fig. 5B, 5C, and Fig. S3D)

Finally, we assessed the functional consequences of impaired ciliogenesis by measuring fluid flow on the embryo surface. Fluorescent beads placed onto control embryos moved along stereotypical trajectories, and automated tracking revealed long, continuous paths with consistent directionality, indicative of coordinated ciliary beating (Fig. 5D, Movie S3). In contrast, bead movement on Cspp1 morphant embryos was disorganized, with frequent pauses and changes in direction that produced intersecting and interrupted trajectories (Fig. 5D, Movie S3). Consistent with this, mean bead speed dropped significantly from ∼175 μm/s in controls to <100 μm/s in morphants (Fig. 5E). Together, these data demonstrate that Cspp1 is required for robust ciliogenesis and effective cilia-driven flow in the *Xenopus* embryonic epidermis.

## Discussion

In this study, we identify CSPP1 as a conserved regulator of vertebrate multiciliogenesis with roles at multiple stages of MCC differentiation. Its dynamic localization across the multiciliogenesis program is consistent with these stage-specific functions. CSPP1 localizes to fibrous granules and deuterosomes during centriole amplification, and to basal bodies and the distal axoneme in mature MCCs. Across mouse airway cultures and *Xenopus* epidermis, loss of CSPP1 disrupts centriole amplification, impairs basal body apical migration, spacing, and rotational polarity, and compromises cilia formation, axoneme structure, and motility. Together, these findings define key cellular functions of CSPP1 in MCCs and provide a framework for understanding how a Joubert syndrome-associated gene can contribute to respiratory phenotypes, including the chronic sinusitis reported in some individuals with CSPP1 mutations [53].

A major strength of our study is the use of two complementary vertebrate MCC models, *in vivo Xenopus* epidermis and *in vitro* mouse airway cultures. These systems share a core multiciliogenesis program but differ in tissue architecture, developmental context, and functional output [31–33]. Comparing them allowed us to distinguish conserved functions from model-specific phenotypes. Across both systems, CSPP1 depletion impaired centriole amplification, ciliogenesis, and ciliary function. This conserved role is further supported by a recent proteomic analysis of an inducible *Xenopus* MCC model, in which CSPP1 was detected from the deuterosome stage through maturation, consistent with a sustained role across multiciliogenesis [34].

Beyond these conserved functions, Cspp1 depletion also impaired MCC intercalation in *Xenopus*. Importantly, centriole amplification and ciliogenesis defects persisted in successfully intercalated cells, supporting a direct requirement for CSPP1 in these processes rather than a secondary consequence of failed intercalation. The intercalation defect resembles that reported after loss of CEP104, a Joubert syndrome protein in the CSPP1 interaction network that destabilizes cytoplasmic microtubules and disrupts MCC intercalation in *Xenopus* [48, 60]. This is consistent with prior work showing that subapical microtubule networks are required for proper MCC integration and organization [44]. Together, these observations suggest that Joubert-associated microtubule regulators may contribute both to axoneme assembly and length control and to the broader microtubule-dependent remodeling events that shape MCC morphogenesis.

Consistent with this broader role, our data place CSPP1 early in the multiciliogenesis pathway, where it contributes to centriole amplification and its coordination with apical remodeling. In *Xenopus*, Cspp1 depletion disrupted deuterosome formation while leaving PCM1-positive fibrous granules intact, positioning CSPP1 downstream of initial fibrous granule assembly. In airway MCCs, CSPP1 loss reduced centriole amplification efficiency and altered the relationship between centriole number and apical surface area, indicating that centriole production was no longer properly matched to apical expansion. In MCCs, dynamic actin networks drive apical surface expansion, whereas the apical microtubule cytoskeleton helps spatially constrain this remodeling and organize basal bodies [44, 61]. In this context, the disrupted apical microtubule network observed upon CSPP1 depletion is consistent with a role for CSPP1 in coordinating centriole amplification with apical remodeling.

This same cytoskeletal disruption likely explains the subsequent defects in basal body integration. Proper basal body apical migration, spacing, and rotational polarity depend on coordinated remodeling of the subapical cytoskeleton [41–44]. Accordingly, the combined spatial defects observed after CSPP1 depletion are consistent with an impaired ability to organize the microtubule network needed to physically anchor and align basal bodies. In the future, defining how CSPP1 couples centriole amplification to apical cortical remodeling may provide insight into the spatiotemporal control of MCC differentiation.

Following apical integration, CSPP1 also contributes to the assembly and maintenance of motile axonemes. Previous work identified CSPP1 at the distal tip of motile cilia and linked it to ciliary length regulation together with CEP104 [48]. Extending these observations, we find that in mature multiciliated cells CSPP1 localizes to basal bodies and the distal axoneme, and that its depletion results in fewer, shorter, and ultrastructurally abnormal cilia with impaired beating. These functions are consistent with recent biochemical studies showing that CSPP1 can stabilize microtubules from the luminal side by capping quiescent ends and repairing damaged lattices [46, 47, 61]. In addition, the similarity between the motile cilia defects observed here and the shortened primary cilia reported in Joubert syndrome-related contexts raises the possibility that a conserved CSPP1-CEP104 module contributes to axoneme length control across distinct ciliary types [62, 63]. The precise biophysical role of CSPP1 in the motile axoneme remains to be defined, including whether it primarily promotes end stabilization, lattice repair, or both.

More broadly, our findings support the emerging clinical paradigm that shared regulatory modules link primary and motile ciliopathies. CSPP1 is particularly informative in this regard, as sensory and respiratory features co-occur within the same individuals [53]. This parallels recent evidence that mutations in the Joubert-associated gene KIAA0586 cause concurrent primary and motile ciliopathies through shared defects in basal body migration [16]. In this context, our work shows that a single microtubule-associated protein can function in both primary and motile cilia, contributing to basal body organization and to axonemal growth and stability. Although loss of CSPP1 produces a shared structural defect, namely shortened and abnormal axonemes, the consequences differ according to the architecture and function of each ciliary type, manifesting as impaired Hedgehog signaling in primary cilia and defective motility in MCCs. Together, these findings support a model in which disruption of a shared cytoskeletal assembly pathway can link primary and motile ciliopathy phenotypes across multiple ciliary tissues.

## Materials and Methods

### Mouse experiments

#### Mouse Husbandry and Primary Mouse Tracheal Epithelial Cell (MTEC) Culture

Mice were maintained under specific pathogen-free standard housing conditions at 20–25 °C and 50% relative humidity on a 12 h light and 12 h dark cycle at the Koç University Animal Research Facility. All animal procedures were conducted in accordance with institutional animal care and use guidelines and were approved by the Koç University Animal Experiments Local Ethics Committee (HADYEK approval no. 2024-44). C57BL/6 background mice aged 8–12 weeks were used. Animals were randomly selected from both sexes in equal proportion.

Primary mouse tracheal epithelial cell (MTEC) cultures were established as previously described [64]. Mice were euthanized by CO₂ exposure. Tracheae were dissected and incubated overnight at 4 °C in 1.5 mg/mL Pronase (Roche, 10165921001) prepared in DMEM/F-12 medium (Sigma, D6421) supplemented with 100 U/mL penicillin and 1 mg/mL streptomycin. Following enzymatic digestion, dissociated cells were seeded onto Primaria plates (BD Biosciences, 353803) and incubated for 4 h at 37 °C in MTEC basic medium (BM). BM consisted of DMEM/F-12 with 15 mM HEPES (Sigma, D6421), 1.5 mM L-glutamine (GlutaMAX, Gibco, 35050), 3.6 mM NaHCO₃ (Gibco, 25080), 10% fetal bovine serum (FBS, Gibco, 10500-064), and 0.25 µg/mL fungizone (Invitrogen, 15290-018) to enrich for epithelial progenitor cells.

Airway epithelial cells were collected, counted, and seeded at 50,000 cells per well onto 50 µg/mL rat-tail collagen (BD, 354236) coated 24-well Transwell filters (Corning, 3470). Cells were cultured in MTEC Plus medium supplied to both apical and basolateral chambers for 5–6 days to allow proliferation and tight junction formation. MTEC Plus medium consisted of BM supplemented with 10 µg/mL insulin (Sigma, I6634), 5 µg/mL transferrin (Sigma, T1147), 25 ng/mL epidermal growth factor (BD Biosciences, 354001), 100 ng/mL cholera toxin (Sigma, C8052), 5% FBS, and 70 µg/mL bovine pituitary extract (Hammond Cell Tech, 1077). MTEC Plus medium was freshly supplemented with 50 nM retinoic acid (Sigma, R2625) and 10 µM ROCK inhibitor (Y-27632, Sigma, Y0503) and refreshed every 2 days until confluency.

Air–liquid interface (ALI) conditions were initiated just prior to full confluency to induce differentiation. After ALI initiation, cells were fed only from the basolateral chamber with MTEC serum-free (SF) medium refreshed every 2 days. SF medium consisted of BM supplemented with 5 µg/mL insulin (Sigma, I6634), 5 µg/mL transferrin (Sigma, T1147), 5 ng/mL epidermal growth factor (BD Biosciences, 354001), 25 ng/mL cholera toxin (Sigma, C8052), 1 mg/mL BSA (Sigma, A2153), and 70 µg/mL bovine pituitary extract (Hammond Cell Tech, 1077). SF medium was supplemented with 50 nM retinoic acid. To promote multiciliated cell differentiation, 5 µM DAPT (Sigma, D5942-5MG) was applied from ALI day 1 through ALI day 5. For microtubule stabilization, MTECs were treated at ALI day 2 with 10 µM Taxol for 24 h and fixed at ALI day 3 with ice-cold methanol.

### Cell Lines

Inner medullary collecting duct cells (mIMCD-3, ATCC, CRL-2123) were cultured in DMEM/F-12 50/50 (Pan Biotech, P04-41250) supplemented with 10% FBS (Gibco). Human embryonic kidney cells (HEK293T, ATCC, CRL-3216) were grown in DMEM (Pan Biotech, P04-03590) supplemented with 10% FBS (Gibco). All cultures were maintained at 37 °C in 5% CO₂.

### Plasmids, Transfection, Lentivirus Production, and Transduction

Mouse CSPP1 shRNA targeting the C-terminal region (nucleotides 2585–2606, sequence 5′-GAGGAACCCAATGGATATATT-3′) cloned into the pLKO.1-TRC vector (Invitrogen, The RNAi Consortium) was used for gene silencing. A Mission Scramble pLKO.1 construct (Invitrogen) served as a non-targeting control. Human CSPP1 cDNA was amplified from GFP-CSPP1 plasmid (gift from S.Patzke) and cloned into pCDH-EF1-mNeonGreen-T2A-Puro plasmid (Clontech). Recombinant lentiviruses were produced in HEK293T cells by co-transfection of the shRNA transfer vector or pCDH-mNG-CSPP1 vector with the packaging plasmid pCMVΔR8.74 and the envelope plasmid pMD2.G (VSV-G). Transfections were performed using 25 kDa polyethylenimine (PEI, Sigma-Aldrich) prepared as a 1 µg/µL stock. Viral supernatants were collected 48 h after transfection and used for infections.

For mIMCD-3 transduction, cells were seeded at low confluency (1 × 10⁵ cells per well in 6-well plates) 24 h prior to infection. Cells were incubated with 300 µL lentiviral supernatant (titer not determined). Medium was replaced after 24 h. At 48 h post-infection, cells were passaged and selected with puromycin (3 µg/mL) for 4 days until non-infected control cells were eliminated. For MTEC transduction, 50,000 cells were seeded onto 24-well Transwell filters and transduced with 50 µL lentiviral supernatant. After 48 h, medium was replaced with MTEC Plus medium containing puromycin (2 µg/mL). Selection was maintained for 4 days. Cells were then cultured in MTEC Plus medium, refreshed every 2 days, until confluency.

### Immunoblotting

Virus-infected and puromycin-selected mIMCD-3 cells were washed twice with PBS and lysed in lysis buffer containing 50 mM Tris-HCl (pH 7.6), 150 mM NaCl, 1% Triton X-100, and protease inhibitors. Lysates were tumbled for 20 min at 4 °C and clarified by centrifugation at 15,000g for 15 min at 4 °C. Protein concentrations were determined using Bradford reagent (Bio-Rad). Samples were mixed with 6× Laemmli sample buffer and boiled at 95 °C for 10 min.

Equal amounts of protein were resolved by SDS-PAGE and transferred to nitrocellulose membranes (Bio-Rad). Membranes were blocked in 5% milk in TBST for 1 h at room temperature and incubated with primary antibodies diluted in blocking solution overnight at 4 °C. Membranes were washed extensively in TBST and incubated with secondary antibodies for 1 h at room temperature. Signals were detected on a LI-COR Odyssey Infrared Imaging System.

Primary antibodies used for immunoblotting were rabbit anti-CSPP1 (1:1000, Proteintech 11931-1-AP) and mouse anti-γ-tubulin (1:1000, Sigma-Aldrich T6557). Secondary antibodies were IRDye 680 and IRDye 800 conjugates used at 1:10,000 (LI-COR Biosciences, 926-68070 and 926-32211). Band intensities were quantified in ImageJ. Background-subtracted CSPP1 signals were normalized to γ-tubulin and then normalized to 1. Plots were generated in GraphPad Prism 10.

### Immunofluorescence of MTECs, trachea, and ependyma, antibodies, and microscopy

For MTEC immunofluorescence, cells grown on 24-well Transwell filters were rinsed twice with PBS and fixed in ice-cold methanol at −20°C for 10 min. Samples were washed twice in PBS and blocked for 1 h at room temperature in 3% BSA in PBS containing 0.1% Triton X-100. Filters were excised from the plastic support and cut into quarters to allow staining with different antibody combinations. Samples were incubated with primary antibodies diluted in blocking solution overnight at 4°C, washed twice in PBS, and then incubated with Alexa Fluor–conjugated secondary antibodies (1:2000) and DAPI in blocking solution for 1 h at room temperature. After PBS washes, samples were mounted in Mowiol containing N-propyl gallate (Sigma-Aldrich).

For in situ trachea staining, whole tracheae were dissected and fixed in ice-cold methanol at −20°C for 10 min. Subsequent staining steps were performed as described for MTECs.

For ependymal immunofluorescence, lateral wall whole mounts were dissected from postnatal day 4 mouse brains and immunostained using a standard whole-mount protocol as described previously [41].

Primary antibodies used for MTEC, trachea, and ependyma immunofluorescence were mouse anti-Centrin (clone 20H5, 1:500, EMD Millipore 04-1624), rabbit anti-Cep164 (1:1000, Proteintech 22227-1-AP), mouse anti-acetylated α-tubulin (clone 6-11B0-1, 1:2000, Sigma T6793), rabbit anti-DEUP1 (1:200, Sigma HPA010986), rat anti-ZO-1 (1:1000, Santa Cruz sc-33725), mouse anti-γ-tubulin (GTU-88, 1:1000, Sigma-Aldrich T6557), rabbit anti-CSPP1 (1:50, Proteintech 11931-1-AP), mouse anti-ODF2 (1:200, Novus/Abnova H00004957-M01), mouse anti-PCM1 (G6 clone, 1:500, Santa Cruz sc-398365), and mouse anti-α-tubulin (DM1A, 1:500, Sigma T926).

Confocal imaging of MTECs and trachea was performed on Leica TCS SP8 and Leica Stellaris 8 inverted laser-scanning confocal microscopes equipped with HC PL APO CS2 oil-immersion objectives (63×/1.4 NA or 40×/1.3 NA). The pinhole was set to 1 Airy unit. Z-stacks were acquired as 15–30 optical sections with 0.2–0.5 µm step size using line averaging (2–3×) and unidirectional scanning at 400 Hz. Images were collected at 512 × 512 or 1024 × 1024 pixel resolution with optical zoom as appropriate. Acquisition and primary processing were performed in Leica Application Suite X (LAS X). Deconvolution was performed using LAS X Lightning or Huygens Professional (Scientific Volume Imaging), as indicated.

Confocal imaging of ependymal whole mounts was performed on a Leica SP8 laser-scanning confocal microscope equipped with a 63×/1.4 NA oil-immersion objective. Z-series were acquired at 0.3 µm step size.

Three-dimensional structured illumination microscopy (3D-SIM) was performed on an Elyra 7 system with Lattice SIM² (Carl Zeiss Microscopy) using a Plan-Apochromat 63×/1.4 NA oil objective. Excitation was provided by 488, 561, and 633 nm laser lines. Z-series were acquired at 110 nm axial intervals across 5–10 µm volumes using an sCMOS camera (pco.edge 4.2 CL HS). Raw SIM datasets were reconstructed in ZEN Black. Channel registration was calibrated using super-resolution TetraSpeck bead standards (Zeiss).

### Image analysis of MTEC datasets

*Basal body and cilia correlation analysis:* 3D-SIM datasets were used to resolve individual basal bodies. Basal body number was quantified using the TrackMate plugin in Fiji/ImageJ. Centrin and ZO-1 channels were separated and maximum intensity projections were generated. Cell boundaries were defined from the ZO-1 channel by manually drawing a region of interest for each cell. TrackMate was run within each cell region of interest using an estimated object diameter of 0.30 µm and a quality threshold of 800. False-positive detections were removed by manual curation. TrackMate outputs included basal body counts and x and y coordinates. Per-cell outputs were exported to Excel and analyzed in GraphPad Prism 10.

*Basal body distribution and nearest neighbor index:* Basal body coordinates exported from TrackMate were saved as .csv files for each cell. A Python script was used to calculate dot number, mean nearest-neighbor distance derived from dot coordinates, and the expected nearest-neighbor distance under complete spatial randomness. The mean nearest-neighbor distance was normalized to the expected value to obtain the nearest neighbor index. Values at or below 1 indicate clustering, and values above 1 indicate increasingly uniform distributions. Outputs were imported into GraphPad Prism 10 for statistical analysis.

*Basal body planar polarity:* Basal body and basal foot orientation were quantified using CEP164 to mark basal bodies and γ-tubulin to mark basal feet. In Fiji/ImageJ, vectors were drawn manually from each basal body toward the corresponding basal foot using the arrow tool. Vector angles were exported and analyzed using Python to generate rose plots and to compute the circular standard deviation for each cell.

### High-speed video microscopy and ciliary beating frequency

Ciliary beating frequency of MTECs at ALI day 8 was measured using high-speed video microscopy and Sisson-Ammons Video Analysis (SAVA) software [65, 66]. Videos were acquired using a Basler acA1300-200um camera mounted on an inverted phase-contrast microscope (Nikon Eclipse TS100) with a 40× dry objective. Images were recorded at 640 × 480 pixels at 120–150 frames per second. Movies were acquired for 1 min with 15 s intervals. CBF was calculated in SAVA using slow-motion and real-time replay of the apical surface.

### Scanning and transmission electron microscopy

MTECs at ALI day 8 on 24-well Transwell filters were fixed in freshly prepared solution containing 2.5% glutaraldehyde (Sigma, 8.20603), 2% paraformaldehyde (pH 7.4), 0.1 M sodium cacodylate, and 1.2 mM CaCl₂, followed by PBS washes for 30 min. Samples were post-fixed in 2% osmium tetroxide (Electron Microscopy Sciences, RT19140) and 1% uranyl acetate (Electron Microscopy Sciences, 22400). Samples were dehydrated through a graded ethanol series (30% to 100%) and acetone (Sigma, 24201). For scanning electron microscopy, samples were critical-point dried (Leica EM CPD300), sputter-coated with gold (Leica EM ACE 200), mounted on aluminum stubs, and imaged on a Zeiss Gemini SEM 500. For transmission electron microscopy, samples were embedded using an epoxy embedding medium kit (Sigma-Aldrich, 45359-1EA-F). Ultrathin sections (Leica UC7) were carried out from the substrate face of the cells, deposited on formvar-coated grids and contrasted by 1% uranyl acetate for 8 minutes and 2,6% lead citrate for 4 minutes. Images were acquired using a Tecnai G2 (Thermofisher, USA), running an LaB6 crystal at 200kV and equipped with a 2K Veleta camera (Olympus, Japan). Cilium length in multiciliated cells was quantified from SEM images using ImageJ.

### *Xenopus* experiments

#### Plasmids, mRNA synthesis, morpholinos, and splice-blocking morpholino validation

Expression vectors were synthesized by EzyVec (France). The *Xenopus laevis* Cspp1 coding sequence was designed based on the Xenbase reference (XB-GENE-5770628). Vectors contained CMV and SP6 promoters and encoded either a morpholino-sensitive Cspp1 coding sequence or a version resistant to the Cspp1 translation-blocking morpholino. The morpholino-resistant sequence was 5′-ATGGGTGAAGAGCTGGAC-3′. Constructs encoded Cspp1 with or without an in-frame mCherry tag. All plasmids were sequence verified.

Capped mRNAs were synthesized from linearized templates using the SP6 mMESSAGE mMACHINE Kit (Ambion Life Technologies) and purified with the MEGAclear Kit (Ambion Life Technologies). Two morpholinos targeting Cspp1.L (XB-GENE-5770628) were designed by GeneTools: a translation-blocking morpholino (5′-GATCTAGTTCCTCCCCCTAC-3′) and a splice-blocking morpholino (5′-TATTGTTGTGTTTACCCTTATTTCC-3′). A fluorescein-tagged splice-blocking morpholino was used for validation and flow analysis experiments.

For splice-blocking morpholino validation, total RNA was purified from animal caps using the RNeasy Kit (Qiagen). PCR was performed using GoTaq G2 Flexi DNA Polymerase (Promega) to detect intron 2 retention, using primers in exon 2 (forward, 5′-GAAGAACAACGGGCCAAACT-3′) and exon 3 (reverse, 5′-CTTGCCTCCTGATTGCTTCC-3′).

### *Xenopus* Embryo Culture and Microinjections

All *Xenopus* experiments were performed in accordance with Directive 2010/63/EU and were approved by the Direction départementale de la Protection des Populations, Pôle Alimentation, Santé Animale, Environnement, des Bouches du Rhône (agreement number G 13055 21). Eggs obtained from NASCO females were fertilized in vitro, dejellied, and cultured using standard protocols [36].

For Cspp1 localization, morpholino-resistant Cspp1-mCherry mRNA (200 pg per blastomere) was injected into a ventral blastomere at Nieuwkoop and Faber stage 3. Embryos were cultured at 18 °C to stage 17 or stage 27–30 and fixed for analysis.

For Cspp1 knockdown, Cspp1 translation-blocking morpholino (10 ng per blastomere) or Cspp1 splice-blocking morpholino (40 ng per blastomere) was injected into the animal-ventral blastomere at stage 4. GFP-GPI (200 pg per blastomere) was co-injected as a tracer. Embryos were cultured at 18 °C to stage 17 or stage 27–30 and fixed.

For rescue experiments, morpholinos were injected into a ventral blastomere at stage 3, followed by injection of morpholino-resistant Cspp1 mRNA into the same blastomere at stage 4. Cspp1 translation-blocking morpholino (10 ng per blastomere) with GFP-GPI (200 pg per blastomere) was rescued with morpholino-resistant Cspp1 mRNA (1000 pg per blastomere). Fluorescein-tagged splice-blocking Cspp1 morpholino (10 ng per blastomere) was rescued with morpholino-resistant Cspp1 mRNA (800 pg per blastomere). Embryos were cultured at 13 °C to stage 27–30 and fixed.

For splice-blocking morpholino validation by RT-PCR, fluorescein-tagged splice-blocking morpholino (10 ng per blastomere) was injected into two blastomeres at stage 2. Animal caps were dissected at stage 10 from uninjected embryos or embryos injected with GFP-GPI control or Cspp1 morpholinos (n = 50 embryos per condition) in 1× MBS and cultured in 0.5× MBS until matched controls reached stage 14. Caps were snap frozen at stage 14 and stored at −80 °C.

### *Xenopus* PCM1 antibody generation and embryo immunofluorescence

Embryos were fixed in 4% paraformaldehyde containing 0.1% Triton X-100 for 30 min. After three washes in PBS, embryos were blocked in 3% BSA in PBS for 1 h at room temperature and incubated with primary antibodies diluted in blocking solution overnight at 4 °C. Primary antibodies were chicken anti-GFP (1:500, AvesLab GFP-1020), rat anti-RFP (1:500, Chromotek 5F8), mouse anti-acetylated tubulin (1:1000, Sigma-Aldrich T7451), mouse anti-Centrin (1:1000, Sigma-Aldrich 04-1624), mouse anti-γ-tubulin (1:500, Abcam ab11316), rabbit anti-Pericentrin (1:5000) [67], rabbit anti-Deup1 (1:800) [34], and rabbit anti-PCM1 (1:5000, this study). Following PBS washes, embryos were incubated with appropriate Alexa Fluor- or Cy3-conjugated secondary antibodies (1:500–1:800) for 1 h at room temperature and mounted in Mowiol.

Rabbit polyclonal antibodies against *Xenopus laevis* PCM1 were generated by immunization with a recombinant N-terminal PCM1 fragment corresponding to residues 1 to 300. Antibody specificity was validated by immunoblotting in COS-1 cells transfected with PCM1 expression constructs using FuGENE HD (Roche). Cells were lysed in modified RIPA buffer, extracts were resolved by SDS-PAGE, and proteins were transferred to Optitran membranes (Whatman) for immunoblot analysis.

### Microscopy and image analysis of *Xenopus* datasets

Confocal images of immunostained embryos were acquired on a Zeiss LSM 880 laser-scanning microscope using a 63×/1.4 NA oil-immersion objective. Z-stacks were processed as maximum-intensity projections and analyzed in Fiji.

PCM1 granule number and size were quantified in Fiji. A 20 µm × 20 µm region of interest was defined for each cell. The PCM1 channel was thresholded, and particles were quantified using the Analyze Particles function. Particle number and size were recorded and compared between GFP-GPI injected control cells and Cspp1 morpholino-injected cells.

For ciliary motility assays, flow at the surface of stage 30 embryos was measured by fluorescent bead tracking. Embryos were anesthetized in 0.02% MS-222 prepared in 0.1× MBS and immobilized on agarose. Ten-micrometer fluorescent microbeads (Thermo Fisher Scientific F8834) were deposited on the dorsal flank. Videos were acquired at 25 frames per second using a Nikon SMZ18 stereomicroscope at 3× magnification equipped with a Hamamatsu ORCA-Fusion camera (C14440) using 10 ms exposure. Movies were analyzed in Fiji using TrackMate. For each movie, a 350 × 350 pixel region of interest was selected in an area containing sufficient bead tracks. Beads were detected using the Difference of Gaussian detector and tracked using the Kalman tracker. Bead velocities were exported for plotting and statistical analysis in GraphPad Prism 10. After imaging, embryos were fixed and immunostained to confirm the corresponding ciliary phenotype.

### Human airway epithelial cell culture, immunostaining, and microscopy

Cryopreserved human bronchial epithelial cells (HBEpC, C-12640, PromoCell, maximum passage 3) were thawed in T75 flasks in complete PneumaCult-Ex Plus medium and cultured at 37°C with 5% CO2. Medium was changed 24 h after thawing and every two days thereafter until cells reached 50–60% confluence. After trypsinization, cells were seeded at 250,000 cells/ml onto Transwells (Corning 6.5 mm Transwell inserts with 0.4 µm pore polyester membranes, no. 3470, Costar), with both apical and basal chambers filled with complete PneumaCult-Ex Plus medium. Just before reaching confluence, cultures were transferred to air–liquid interface conditions by removing medium from the apical chamber and replacing the basal medium with complete PneumaCult-ALI medium. Basal medium was changed every two days.

ALI day 27 human airway epithelial cultures were fixed in methanol for 6 min and washed three times in PBS containing 0.1% Tween-20. Samples were blocked in 3% BSA in PBS for 15 min at room temperature and incubated with rabbit anti-CSPP1 (1:200, Proteintech 11931-1-AP), diluted in PBS containing 3% BSA for 45 min at room temperature. After two washes in PBS containing 0.1% Tween-20, samples were incubated with Alexa Fluor-coupled anti rabbit secondary antibodies diluted in PBS containing 3% BSA for 30 min at room temperature. After two additional washes in PBS containing 0.1% Tween-20, Transwell membranes were excised and mounted in Mowiol between slides and coverslips. Confocal images were acquired as Z-series with 0.3 µm step size on a Leica SP8 laser-scanning microscope using a 63×/1.4 NA oil-immersion objective.

### Statistical analysis

All statistical analyses were performed in GraphPad Prism 10 (GraphPad Software). Data normality was assessed using the D’Agostino and Pearson test when n exceeded 50, or the Shapiro–Wilk test when n was below 50. For comparisons between two groups, an unpaired two-tailed Student’s t test was used for normally distributed data and the Mann–Whitney test was used for non-normally distributed data. For comparisons of more than two groups, ordinary one-way ANOVA was used. For assay-specific analyses, including proportion-based outcomes and particle or flow measurements, the statistical tests, definition of n, and multiple-comparison corrections are indicated in the corresponding figure legends. Error bars represent standard deviation. Statistical significance is indicated as ns for P greater than 0.05, * for P up to 0.05, ** for P up to 0.01, *** for P up to 0.001, and **** for P up to 0.0001.

## Supporting information

AllSupp

## Acknowledgements

This work received support from the French government under the France 2030 investment plan, as part of the Initiative d’Excellence d’Aix-Marseille Université AMIDEX (AMX-21-PEP-042, “Pépinière d’Excellence 2021” to Ca.Bo.), Cancéropôle PACA–GEFLUC (Programme Emergence 2021 to Ca.Bo.), the French National Research Agency (ANR-22-CE13-0027, PHACIL to Ca.Bo.) and by the French Ministry for Europe and Foreign Affairs (MEAE) and the Ministry of Higher Education, Research, and Innovation (MESRI) (PHC Bosphore 2022, 47980WA to Ca.Bo and JD). Cl.Bo. is supported by a PhD fellowship from the Ligue Nationale Contre le Cancer. The IBDM photonic and electron microscopy imaging platforms are members of the national infrastructure France-BioImaging (PICsL-FBI) (https://ror.org/01y7vt929), supported by the French National Research Agency (ANR-24-INBS-0005, FBI BIOGEN), and are also part of the Marseille Imaging Institute, an Excellence Initiative of Aix-Marseille Université AMIDEX within the French “Investissements d’Avenir” programme (AMX-19-IET-002). Authors thank Emmanuelle Leblanc and Florian Roguet for *Xenopus* husbandry and Nilhan Coşkun for mouse husbandry. This work was also supported by the International Centre for Genetic Engineering and Biotechnology (ICGEB; CRP/TUR21-01 to E.N.F.K.), the European Research Council (ERC) under the European Union’s Horizon Europe programme (grant agreement 101078097 to E.N.F.K.), and TÜBİTAK ARDEB 1001 (grant 123Z442 to E.N.F.K.).

## Author contribution

Conceptualization: E.N.F.K. and Ca.Bo.; Investigation: I.S.D., Cl.Bo., J.D., M.T., M.S.A., E.Y., O.K., H.B., V.T., O.R., N.B., and Ca.Bo.; Formal analysis: I.S.D., Cl.Bo., J.D., Ca.Bo., and E.N.F.K.; Writing: E.N.F.K. and Ca.Bo.; Editing: all authors; Funding acquisition: E.N.F.K. and Ca.Bo.; Supervision: E.N.F.K. and Ca.Bo.

