## Supplementary material for "The Joubert syndrome protein CSPP1 is a conserved regulator of vertebrate multiciliogenesis and motile cilia function": AllSupp

### Figure S1

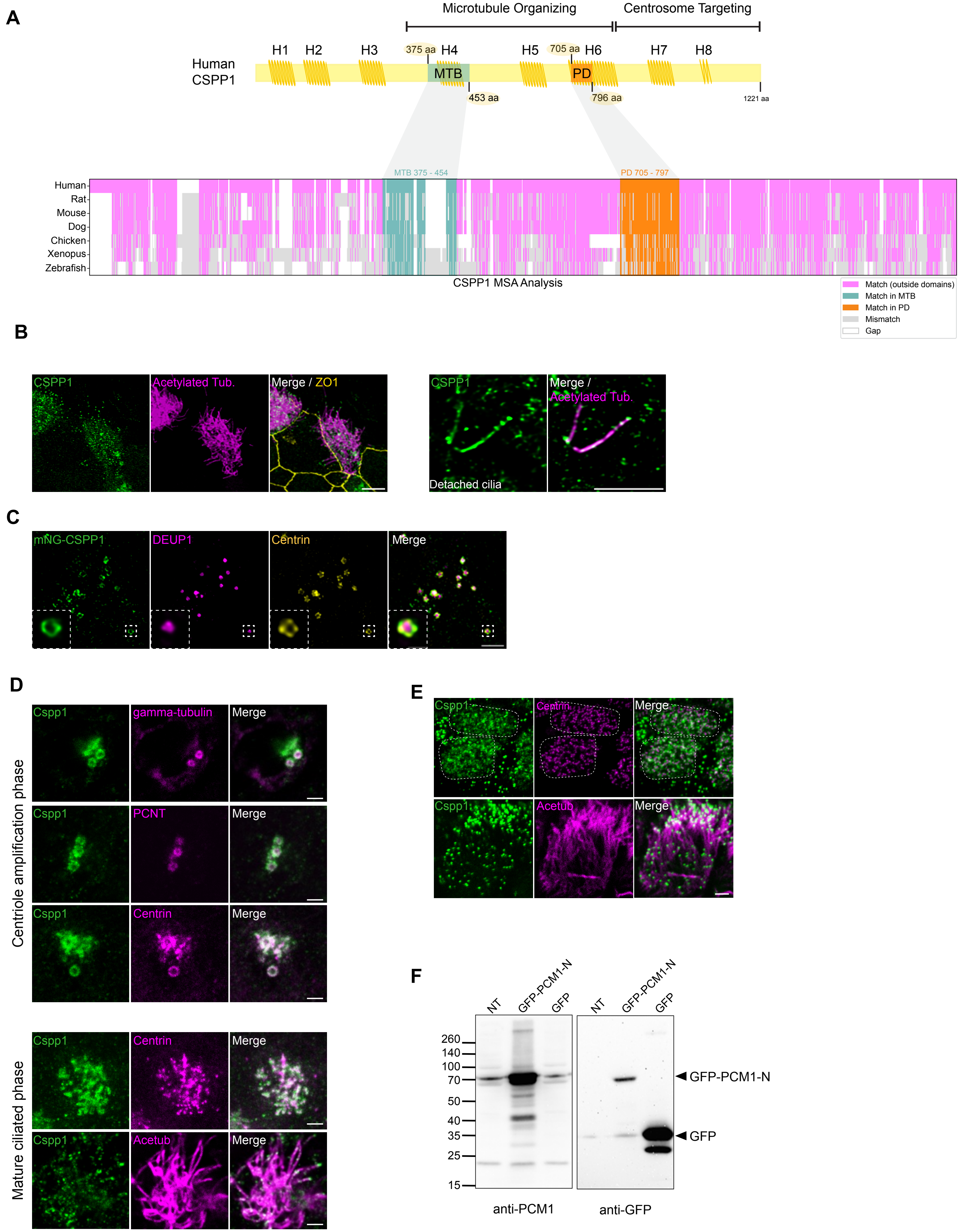
A
Microtubule Organizing
Centrosome Targeting
Human CSPP1
H1
H2
H3
375 aa
H4
H5
705 aa
H6
H7
H8
MTB
PD
453 aa
796 aa
1221 aa
MTB 375 - 454
PD 705 - 797
Human
Rat
Mouse
Dog
Chicken
Xenopus
Zebrafish
CSPP1 MSA Analysis
Match (outside domains)
Match in MTB
Match in PD
Mismatch
Gap
B
CSPP1
Acetylated Tub.
Merge / ZO1
CSPP1
Merge / Acetylated Tub.
Detached cilia
C
mNG-CSPP1
DEUP1
Centrin
Merge
D
Centriole amplification phase
Cspp1
gamma-tubulin
Merge
Cspp1
PCNT
Merge
Cspp1
Centrin
Merge
Mature ciliated phase
Cspp1
Centrin
Merge
Cspp1
Acetub
Merge
E
Cspp1
Centrin
Merge
Cspp1
Acetub
Merge
F
NT
GFP-PCM1-N
GFP
NT
GFP-PCM1-N
GFP
260
140
100
70
50
40
35
25
15
GFP-PCM1-N
GFP
anti-PCM1
anti-GFP

**Figure S1: Structural conservation of CSPP1 and validation of reagents and its localization in diverse multiciliated tissues.**

**(A) CSPP1 domain organization and conservation. Top:** Schematic representation of human CSPP1 protein structure, indicating the eigth helical domains (H1–H8), the microtubule-binding domain (MTB), and pausing domain (PD). **Bottom:** Multiple Sequence Analysis (MSA) plot showing amino acid conservation across vertebrates (Human, Rat, Mouse, Dog, Chicken, *Xenopus*, Zebrafish). The central MTB and PD regions show high conservation.

**(B) CSPP1 localization to basal bodies and cilia in *ex vivo* trachea. Left:** Maximum intensity projection of mouse tracheal tissue stained for CSPP1 (green), Acetylated Tubulin (magenta; cilia), and ZO-1 (yellow; tight junctions). **Far Right:**"Detached cilia" inset showing punctate CSPP1 localization along the ciliary axoneme and enrichment at the tip. **Scale bars:** 5 µm.

**(C) Recruitment of exogenous CSPP1 to deuterosomes.** Representative image of MTECs expressing mNeonGreen-CSPP1 (mNG-CSPP1, green). mNG-CSPP1 colocalizes with the deuterosome marker DEUP1 (magenta) and Centrin (yellow) during centriole amplification. **Scale bars:** 2 µm. Inset is 4X zoom.

**(D) CSPP1 localization in mouse ependymal cells.** Representative immunofluorescence images of mouse ependymal cells differentiating *in vivo*. **Centriole amplification phase:** During early stages, CSPP1 (green) associates with deuterosomes and nascent centrioles, colocalizing with gamma-tubulin, PCNT, and Centrin (magenta). **Mature ciliated phase:** In fully differentiated ependymal cells, CSPP1 (green) localizes to basal bodies marked by Centrin (magenta) and ciliary axonemes marked by Acetylated Tubulin (magenta). **Scale bars:** 2 µm.

**(E) CSPP1 localization in human airway epithelial cultures.** CSPP1 (green) localizes to basal bodies marked by Centrin (magenta) and ciliary axonemes marked by Acetylated Tubulin (magenta). The apical cell surface is outlined with dashed lines. **Scale bars:** 2 µm.

**(F) Validation of the *Xenopus* anti-PCM1 antibody.** Western blot analysis of lysates from cells expressing GFP-tagged *Xenopus* PCM1 N-terminus (1-300) or GFP control, alongside non-transfected (NT) controls. The custom anti-PCM1 antibody (left blot) specifically detects the GFP-PCM1-N (1-300) fusion protein at the expected molecular weight, matching the anti-GFP control blot (right).

### Figure S2

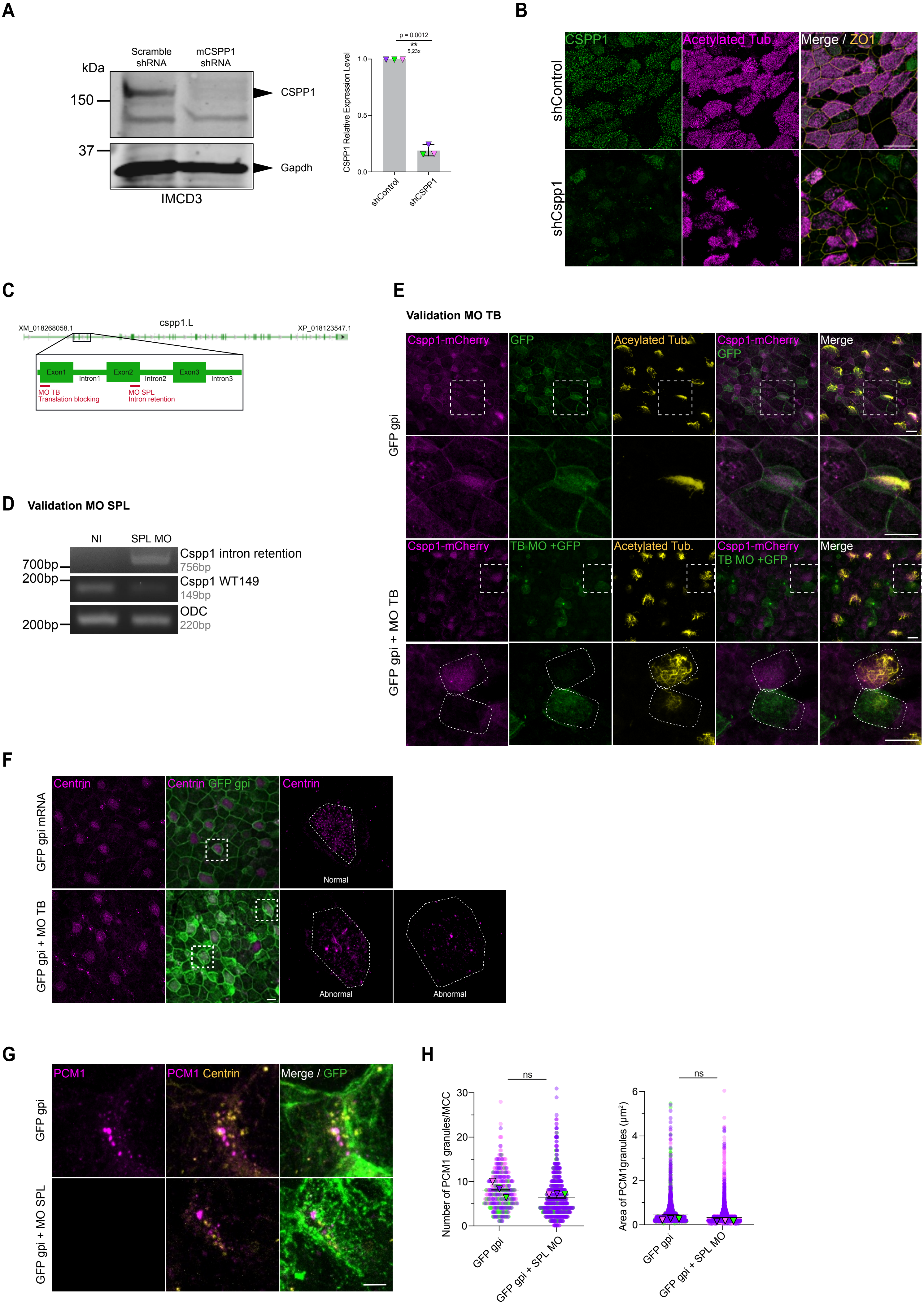
A
Scramble shRNA
mCSPP1 shRNA
kDa
150
37
CSPP1
Gapdh
IMCD3
p = 0.0012
**
5,23x
CSPP1 Relative Expression Level
shControl
shCSPP1
B
CSPP1
Acetylated Tub.
Merge / ZO1
shControl
shCspp1
C
XM_018268058.1
cspp1.L
XP_018123547.1
Exon1
Intron1
Exon2
Intron2
Exon3
Intron3
MO TB
Translation blocking
MO SPL
Intron retention
D
Validation MO SPL
NI
SPL MO
700bp
200bp
200bp
Cspp1 intron retention
756bp
Cspp1 WT149
149bp
ODC
220bp
E
Validation MO TB
Cspp1-mCherry
GFP
Aceylated Tub.
Merge
TB MO +GFP
Acetylated Tub.
GFP gpi
GFP gpi + MO TB
F
Centrin
Centrin GFP gpi
Normal
Abnormal
GFP gpi mRNA
GFP gpi + MO TB
G
PCM1
PCM1 Centrin
Merge / GFP
GFP gpi
GFP gpi + MO SPL
H
ns
Number of PCM1 granules/MCC
Area of PCM1granules (µm²)
GFP gpi
GFP gpi + SPL MO

**Figure S2. Validation of *Cspp1* depletion reagents and analysis of fibrous granules**

**(A) Validation of CSPP1 knockdown in IMCD3 cells.** IMCD3 cells were transduced with Scramble (shControl) or *Cspp1* (shCSPP1) shRNA lentiviruses and selected with puromycin. (Left) Representative Western blot analysis of cell lysates immunoblotted for CSPP1 (~150 kDa) and GAPDH (loading control). (Right) Quantification of relative CSPP1 protein levels. *Cspp1* shRNA significantly reduces protein expression. Data represent mean ± SD from three biological replicates (n=3). Statistical significance determined by paired, two-tailed Student's t-test. **P<0.01.

**(B) Validation of *Cspp1* depletion in MTECs.** Representative immunofluorescence images of MTECs transduced with the indicated lentiviruses, fixed at ALI12, and stained for CSPP1 (green), Acetylated Tubulin (magenta), and ZO-1 (yellow). *Cspp1* depletion results in the loss of specific CSPP1 signal at the apical surface. Scale bar: 20 μm.

**(C) Schematic of *Xenopus laevis cspp1.L* gene structure.** Diagram showing the exon-intron organization of *cspp1.L*(Xenbase: GENE-5770628). The target binding sites for the Translation-Blocking (TB MO, exon 1) and Splice-Blocking (SPL MO, exon 2-intron 2 junction) morpholinos are indicated in red.

**(D) Validation of the *Cspp1* SPL MO by RT-PCR.** Total RNA was extracted from animal caps of non-injected (NI) embryos or embryos injected with the *Cspp1* Splice-Blocking morpholino (SPL MO) at the 2-cell stage. RT-PCR analysis confirms the loss of the wild-type *cspp1* transcript (149 bp) and the appearance of a larger intron-retention product (756 bp) in morphants. ODC serves as a loading control.

**(E) Validation of the Cspp1 TB MO efficiency in vivo.** Representative confocal images of stage 30 (st30) Xenopus laevis embryonic epidermis. Embryos were injected at stage 3 with MO-sensitive Cspp1-mCherry mRNA, followed by injection at stage 4 with either GFP-GPI mRNA alone (top row) or GFP-GPI mRNA together with the *Cspp1* TB morpholino (bottom row). Cells were stained for GFP-GPI (green), Cspp1-mCherry (magenta), and Acetylated Tubulin (yellow). The TB MO effectively suppresses the translation of the reporter, evidenced by the loss of Cspp1-mCherry signal in GFP-positive cells. Scale bar: 20 μm. Inset is 3X zoom.

Higher magnification images show that control cells (GFP-GPI only) display apical Cspp1 localization and a dense ciliary layer, whereas GFP-positive cells co-injected with TB morpholino exhibit loss of Cspp1 signal and severe ciliary defects. Neighboring non-injected cells retain normal Cspp1 localization and cilia density.

**(F) Centriole amplification defects following *Cspp1* TB MO injection.** Representative confocal images of stage 30 *Xenopus laevis* embryonic epidermis injected with GFP mRNA alone (Control) or co-injected with the *Cspp1* TB MO. Cells were stained for the lineage tracer GFP (green) and Centrin (magenta). While control cells show a normal, homogeneous distribution of basal bodies, TB morphants display abnormal centriole numbers and distributions characterized by aggregates and centrin filament formation. Similar results were obtained from three biological replicates. Scale bar: 20 μm. (Quantification is provided in main Figure 2). Inset is 5X zoom. The apical cell surface is outlined with dashed lines.

**(G) Analysis of Fibrous Granules at stage 17.** Representative confocal images of *Xenopus* epidermal MCCs injected with the indicated constructs and stained for PCM1 (magenta, fibrous granules), Centrin (yellow), and GFP (lineage tracer). Scale bar: 5 µm.

**(H) Quantification of Fibrous Granule number and size.** The number of PCM1 particles per MCC was quantified from images as in (G). No significant difference was observed between control and *Cspp1* morphants. Data represent mean ± SD from three biological replicates (n > 400 cells analyzed from >9 embryos per condition). Statistical significance determined by Mann-Whitney test. The mean surface area ($\mu m^2$) of PCM1 granules was quantified. *Cspp1* depletion does not alter fibrous granule size. Data represent mean ± SD from three biological replicates (n > 400 cells analyzed from >9 embryos per condition). Statistical significance determined by Mann-Whitney test.

*P<0.05; **P<0.01, ***P<0.001, ****P<0.0001; ns, not significant.

### Figure S3

**A**

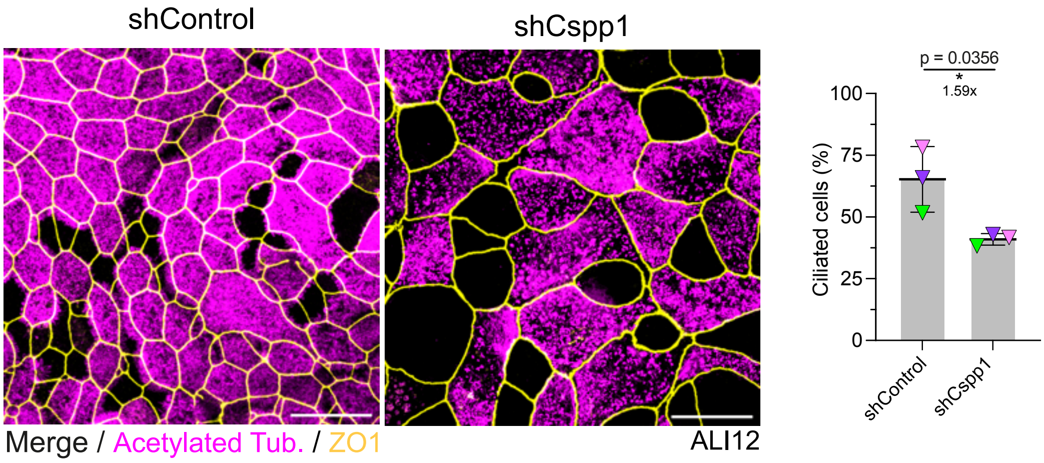


**B** **SUPP DATA** MO TB

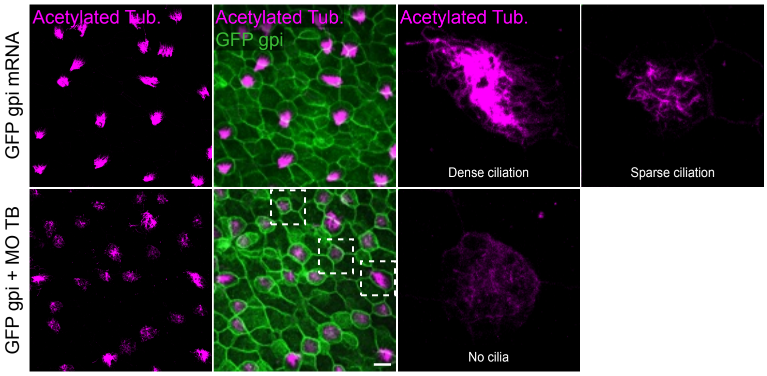


**C** **MO-Res Cspp1 mRNA toxicity test**

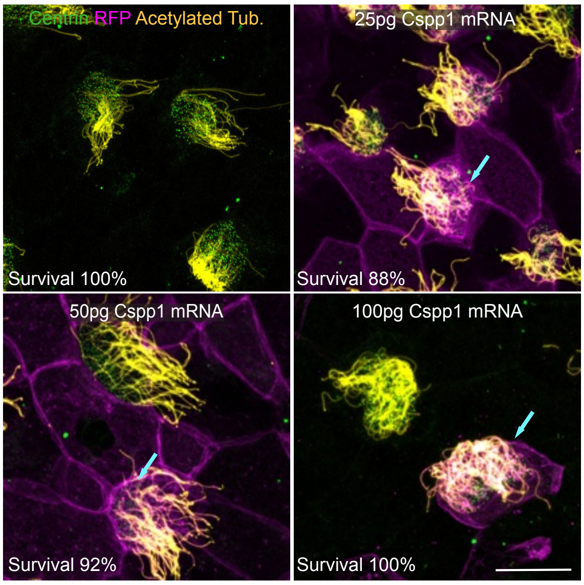


**D** **Rescue**

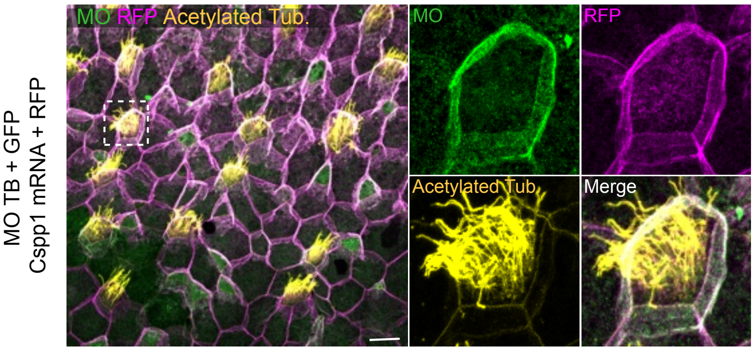

**Figure S3. Characterization of motile cilia formation defects in MTECs and *Xenopus* embryos**

**(A) Percentage of ciliated cell population at ALI12 in MTECs.** The percentage of the ciliated cell population was quantified from the late (ALI12) stage of MTECs using ImageJ. Representative images show cells stained for Acetylated Tubulin (magenta) and ZO-1 (yellow). Graph: *Cspp1* depletion significantly reduces the percentage of multiciliated cells (MCCs). Results are from three biological replicates; colored triangles represent different replicates. Statistical significance was determined by paired, two-tailed Student's t-test. *P<0.05. Scale bar: 20 μm.

**(B) Ciliogenesis defects following *Cspp1* TB MO injection.** Representative confocal images of stage 30 *Xenopus laevis* embryonic epidermis injected with GFP-GPI mRNA alone (top row) or co-injected with the *Cspp1* TB MO (bottom row). Cells were stained for GFP-GPI (green) and Acetylated Tubulin (magenta). Cilia distribution profiles are defined as normal (dense, evenly distributed cilia of uniform length) or abnormal (sparse, short, or absent cilia). Similar results were obtained from three biological replicates. Scale bar: 20 μm. (Quantification is reported in **Main Figure 5B**). Inset is 5X zoom.

**(C) Toxicity test of MO-resistant *Cspp1* mRNA.** Representative confocal images of stage 30 *Xenopus laevis*multiciliated cells either non-injected or co-injected with the indicated amounts (25, 50, or 100 pg) of *Cspp1* mRNA together with 200 pg of RFP mRNA. Cells were stained for RFP (magenta, membrane tracer), Acetylated Tubulin (yellow), and Centrin (green). At stage 30, both ciliogenesis and centriole biogenesis were unaffected following injection of increasing doses of *Cspp1* mRNA. Embryo survival rates were comparable across conditions (percentages indicated on panels). Scale bar: 20 μm.

**(D) Rescue of ciliogenesis defect in TB morphants by *Cspp1* reintroduction.** Representative confocal images of stage 30 *Xenopus laevis* embryonic epidermis injected at stage 3 with the *Cspp1* TB morpholino (MO) and GFP (green), followed by co-injection of MO-resistant *Cspp1* mRNA and membrane-targeted RFP (magenta). Cells were stained for GFP (green), RFP (magenta), and Acetylated Tubulin (yellow). Cilia distribution profiles were categorized as normal (evenly distributed cilia of uniform length) or abnormal. The rescue construct restores normal ciliation in morphant cells. Similar results were obtained in three independent biological replicates (n=3). Scale bar: 20 μm. (Quantification is presented in **Main Figure 5B**). Inset is a 3X zoom.

*P<0.05; **P<0.01, ***P<0.001, ****P<0.0001; ns, not significant.
